# Can Hypnotic Susceptibility be Explained by Bifactor Models? Structural Equation Modeling of the Harvard Group Scale of Hypnotic Susceptibility - Form A

**DOI:** 10.1101/2021.04.29.441926

**Authors:** Anoushiravan Zahedi, Werner Sommer

**Affiliations:** Department of Psychology, Humboldt-Universitat zu Berlin, Rudower Chaussee 18, 12489, Berlin, Germany; German Institute of Human Nutrition, Decision Neuroscience and Nutrition, Arthur-Scheunert-Allee 114-116, 14558, Nuthetal, Germany; Neuroscience Research Center, Charité-Universitätsmedizin Berlin, Germany; Department of Psychology, Zhejiang Normal University, Jin Hua, China

**Keywords:** Confirmatory factor analysis, Structural equation modeling, Hypnosis, Suggestibility, Hypnotizability, HGSHS:A

## Abstract

Individuals differ in their responsiveness to hypnotic suggestions. However, defining and measuring hypnotizability is contentious because standardized scales, such as the Harvard group scale (HGSHS:A), measure a mixture of general suggestibility and its alteration due to hypnotic induction (hypnotizability). Exploratory factor analyses (FA) of standardized scales indicated their multidimensionality; however, the number and nature of latent factors are debated. We applied Confirmatory FA to the HGSHS:A scores of 477 volunteers and tested several theory-driven models. Scores were best explained by a bifactor model consisting of a G-factor and three correlated minor factors. The presented bifactor model shows that two sources of variability affect HGSHS:A simultaneously. Structural equation modeling revealed that the challenge-ideomotor factor predicts the other two minor factors, implying these suggestions might require more fundamental processes than other types. These results demonstrate the multifaceted and bifactorial structure of hypnotic suggestibility and underscore the desideratum for developing more differentiated scales.

## 1. Introduction

Historically, hypnosis has been explained by two main accounts. The state approach defines hypnosis as an altered state of consciousness similar to yoga or meditation (Elkins et al., 2015). In contrast, the socio-cognitive account (Kirsch & Lynn, 1998; Lynn & Green, 2011; Lynn et al., 1990) emphasizes cognitive, social, and psychological variables involved in responding to hypnotic suggestions (Spanos, 1971; Spanos et al., 1985). As discussed by Jensen et al. (2015) and Lynn and Green (2011), contemporary theories of hypnosis only partially align with these traditional alternative views (for review, see Zahedi & Sommer, 2021), and no consensus has been reached. Independent of ongoing disputes, theories of hypnosis agree on the existence of substantial within- and between-subject variability in responding to hypnotic and posthypnotic suggestions (Shor & Orne, 1963). However, defining and measuring hypnotizability is a more contentious issue. Two approaches can be distinguished.

First, based on studies (e.g., Braffman & Kirsch, 1999; e.g., Mazzoni et al., 2009; McGeown et al., 2012; Parris & Dienes, 2013) that have shown strong correlations between responding to suggestions inside and outside of hypnosis (r = .67 for behavioral scores; r = .82 for subjective scores; Braffman & Kirsch, 1999), Kirsch (1997) concluded that suggestibility and hypnotizability should be separated. He distinguished (I) *suggestibility* as the capability to respond to suggestions regardless of hypnosis, (II) *hypnotic suggestibility* as the capability to respond to suggestions under the influence of hypnosis, and finally, (III) *hypnotizability* as the alteration in suggestibility due to induction of hypnosis (i.e., the difference between hypnotic suggestibility and suggestibility). Unfortunately, as yet, this sophisticated definition of hypnotizability has not been translated into a reliable and valid hypnotizability scale. Noticeably, the idea that suggestibility and hypnotizability can be distinguished is not debuted by Kirsch (1997) and was already discussed by Hilgard (1965b) and Hilgard (1973).

Secondly and more traditionally, one might define hypnotizability as what standardized scales of hypnotic susceptibility are measuring. Noteworthy, in this definition, hypnotizability is equated with hypnotic suggestibility in terms of Kirsch’s (1997) description of hypnotizability. Two of the most commonly employed hypnotic-susceptibility scales are the Stanford scale of hypnotic susceptibility: form C (SSHS:C; Weitzenhoffer & Hilgard, 1962) and the Harvard group scale of hypnotic susceptibility: form A (HGSHS:A; Shor & Orne, 1962; Shor & Orne, 1963). These two scales are similar in nature, except that HGSHS:A is designed for group administration, whereas the SHSS:C is designed for individual-participant administration. In both scales, a range of suggestions is presented consecutively, and in the end, either participants themselves (in HGSHS:A) or administrators (in the SHSS:C) determine how many of the suggestions had been executed. Hence, participants can receive a score between 0-12 in both HGSHS:A and SHSS:C. Based on their hypnotic-susceptibility scores, participants are conventionally categorized as the high-, medium-, and low-hypnotic-susceptibles. For instance, in HGSHS:A, participants with *scores* ≥ 9, 8 and 4 ≥ *scores* are considered high-, medium-, and low-hypnotic-susceptible, respectively. Both scales are very stable over time; for instance, for the HGSHS:A scores, stability coefficients of .82 (15 – *year retest*) and .71 (25 – *year retest*) have been reported (Piccione et al., 1989). However, the internal consistency of HGSHS:A is at best just acceptable (e.g., Bongartz, 1985; Peter et al., 2014; Robin et al., 2005; Shor & Orne, 1963; Varga et al., 2012). A related concept is the heterogeneity of standardized scales aiming to measure hypnotic susceptibility.

For instance, several previous studies scrutinized the structure of the HGSHS:A scores (e.g., Hilgard, 1965b; McConkey et al., 1980; Oakman & Woody, 1996; Piesbergen & Peter, 2006; Woody et al., 2005) and SSHS:C scores (e.g., Hilgard, 1965b; Woody et al., 2005) by conducting exploratory factor analyses (EFA). Based on these studies, a strong consensus has been reached that the HGSHS:A items do not represent only a single factor. However, there is less consensus about the number and nature of latent factors involved. For instance, even though some studies favor a two-factor model (e.g., Sadler & Woody, 2004), some others prefer a three-factor (McConkey et al., 1980) or even four-factor models (e.g., Woody et al., 2005). Notably, some of the observed differences can be attributed to using different suggestion pools. There is a tentative consensus that the HGSHS:A items measure at least three latent factors (Tab. 1). The first latent factor is characterized by direct-ideomotor suggestions, such as “soon after thinking of your head falling forward, you feel a tendency to make the movement.” The second factor consists of so-called challenge-ideomotor suggestions, such as “your hands feel heavy… too heavy to be lifted”. And the third factor is related to perceptual-cognitive suggestions, such as “you will be increasingly aware of a fly that is going round and round about your head.” However, the third factor is very loosely defined, and in EFA, sometimes only one suggestion is loading on this factor (e.g., McConkey et al., 1980).

**Table 1.** Categorization of the suggestions in HGSHS:A into three proposed minor factors and the results of a seminal exploratory factor analysis (EFA) conducted by McConkey et al. (1980)

| Suggestions | Proposed Specific Factor | EFA <sup>a</sup> |
| --- | --- | --- |
| 1. HEAD FALLING | F1 | Direct Ideomotor |
| 2. EYE CLOSURE | F1 | Direct Ideomotor |
| 3. HAND LOWERING (LEFT HAND) | F1 | Direct Ideomotor |
| 4. ARM IMMOBILIZATION (RIGHT ARM) | F2 | Challenge Ideomotor |
| 5. FINGER LOCK | F2 | Challenge Ideomotor |
| 6. ARM RIGIDITY (LEFT) | F2 | Challenge Ideomotor |
| 7. MOVING HANDS TOGETHER | F1 | Direct Ideomotor |
| 8. COMMUNICATION INHIBITION | F2 | Challenge Ideomotor |
| 9. EXPERIENCING OF A FLY | F3 | Cognitive-Perceptual |
| 10. EYE CATALEPSY | F2 | Challenge Ideomotor |
| 11. POSTHYPNOTIC SUGGESTION | F3 | Cognitive-Perceptual |
| 12. HYPNOTIC AMNESIA | F3 | Cognitive-Perceptual |

*Note:* F1: direct-ideomotor suggestions; F2: challenge-ideomotor suggestions; F3: perceptual-cognitive suggestions.

Why is there only a weak consensus about the number and nature of latent factors in HGSHS:A? All previous studies investigating the structure of hypnotizability scales have utilized EFA (e.g., McConkey et al., 1980; Oakman & Woody, 1996; Piesbergen & Peter, 2006). Even though EFA is an essential and necessary step, it has limitations. These limitations can be overcome by confirmatory factor analysis (CFA) and structural equation modeling (SEM) (Coulacoglou & Saklofske, 2017). (1) In EFA, no explicit theory-driven hypotheses are formulated and tested. Therefore, the interpretations of the derived factors are post hoc and may vary across studies (Coulacoglou & Saklofske, 2017; Harrington, 2009). That means, in EFA, the analysis starts by selecting the number of latent factors; which indicator loads on which factor is then determined by using reduced correlational matrices. This process, however, is not fixed and can be altered by using different rotation techniques, such as orthogonal versus oblique rotations. Therefore, the post hoc interpretation of factors, determined by which indicators loading on them, should be treated cautiously. In contrast, CFA is theory-driven, and factors are defined a priori ^1^. Therefore, we hold that the data-driven nature of EFA is the main reason why results from EFA do not converge on the number and theoretical explanations of latent factors. Consequently, the present study will apply CFA and SEM (Coulacoglou & Saklofske, 2017; Harrington, 2009) for investigating the homogeneity and structure of HGSHS:A as a canonical example of hypnotic-suggestibility scales.

Further, (2) Even though EFAs have shown that variance in the items of HGSHS:A must be explained by more than one factor, it is unclear whether a model, which, in addition to specific minor factors, also assumes a general factor (G-factor) of hypnotizability, will explain the data better than a simple multifactor model. Of particular interest for addressing this question is bifactor modeling (Reise, 2012), which can be implemented only in CFA but not EFA. Bifactor models have addressed long-standing questions, from personality psychology (Musek, 2017) to neuroimaging of individual differences (Cooper et al., 2019) and probably most noticeably in psychometric and intelligence research (Eid et al., 2018).

Finally, (3) SEM allows us to explore predictive relationships between latent variables. Thus, SEM of HGSHS:A factors can address whether there is a special minor factor on which other minor factors regress. Using SEM, we will test whether responsiveness to a certain group of suggestions (as a latent variable) can predict how participants respond to other suggestions at the level of latent factors. Predictive pathway testing is only interpretable when hypotheses are theory-driven and formulated a priori, and therefore, only applicable in conjunction with CFA. Summarizing, several important questions, namely, the number and nature of factors, the existence of a bifactorial G-factor, and predictive pathways between latent factors, can be addressed best by conducting CFA and SEM, which is the general aim of the present paper. Noticeably, previous studies had shown that in hierarchical models of hypnotic-susceptibility scores, it is beneficial to assume a second-level G-factor, which is measured by first-level latent factors (for review, see Sadler & Woody, 2021; Woody & McConkey, 2003). However, G-factors in bifactor models are directly measured by indicators and are assumed orthogonal to other latent factors. Therefore, bifactorial G-factors should be interpreted differently than hierarchical G-factors.

A vital prerequisite to formulating relevant hypotheses for CFA is an appropriate theory. Based on the separable categories of suggestions found by EFA, Woody et al. (2005) concluded that there are multiple hypnotizabilities, and the differences between different categories of suggestions cannot be attributed to mere the difficulty of items. Notably, their results were obtained by analyzing both HGSHS:A and SHSS:C simultaneously, which creates a bigger pool of items than using any of these scales alone. Consequently, the most straightforward theoretical interpretation of multiple hypnotizabilities is to assume distinguishable cognitive mechanisms underlying the different categories of suggestions. Hence, for a theory to be considered appropriate, it should assume at least two distinguishable cognitive processes as underlying mechanisms for responding to different suggestions. To the best of our knowledge, at least two theories make such assumptions (for review, see Zahedi & Sommer, 2021). In the following, these theories will be discussed shortly. Notably, here, we are neither presenting an exhaustive review of hypnosis theories (for an in-depth review, see Zahedi & Sommer, 2021) nor evaluating them. Instead, based on the results of existing EFAs, we try to account for the HGSHS:A data by means of CFA based on an appropriate theory.

The first theory is the *unified cognitive theory* (Brown & Oakley, 2004), grounded on the concept of contention scheduling (Norman & Shallice, 1986). Norman and Shallice (1986) assume two separable control systems to be involved in action production, that is, the supervisory attentional system (SAS) and contention scheduling (CS). The SAS will interfere when the existing response repertoire is insufficient for handling a situation or task. In these cases, either a new schema (i.e., a stimulus-response contingency) must be created, or a well-established (prepotent) schema should be inhibited in favor of a less-established schema. In situations that need less cognitive control and can be handled by existing response repertoire, different sets of potential “source schemata” may compete with each other, and the schema that first exceeds a certain activation threshold will be selected by CS (Norman & Shallice, 1986). Brown and Oakley (2004) defined two mutually exclusive styles of responding to hypnotic suggestions, namely, constructive and concentrative. In the concentrative style, the SAS will be disabled or decoupled, and therefore, cannot be used for responding to suggestions. Hence, CS will be the only system in charge of action control. In contrast, in the constructive style, goal-directed imagination, requiring the SAS, will be used for responding to hypnotic suggestions. These two styles, however, are used to characterize two different groups of participants; for instance, “… subjects using a constructive style may report using a range of strategies to create the suggested experience, whereas concentrative responders are more likely to report that the experience simply occurred” (Brown & Oakley, 2004, p. 172). Hence, even though the unified cognitive theory and its two response styles can be used to understand between-subject differences in responding to suggestions, it does not explain within-subject variance. Therefore, one cannot employ the unified cognitive theory for interpreting the existence of multiple hypnotizabilities within individuals. Notably, we only discussed the unified cognitive theory here for its elaborateness; however, several other theories, such as the three-dimensional theory (Barber, 1999) and the theory of Shor (1962), make similar assumptions. These theories assume there are several distinguishable groups of individuals, and each group will exclusively rely on a specific mechanism for responding to suggestions. Hence, even though these theories can explain interindividual differences in responding to suggestions (i.e., the existence of several different high-hypnotic-susceptibles), they cannot explain intraindividual variances, which is required for proposing multiple latent factors.

The second hypnosis theory is the *simulation-adaptation theory of hypnosis* (SATH; Zahedi & Sommer, 2021) which incorporates three concepts. The first concept is cognitive simulation, suggesting that imagining [here, imagination is not restricted to visual imagery and can be auditory, tactile, and so forth] a stimulus has the same effects as perceiving that stimulus (Farah, 1988). The main difference between imagining and perceiving is that the former, in contrast to the latter, is caused by inner thoughts rather than external stimuli (Hesslow, 2002). For instance, imagining a stimulus not only activates the same brain areas but also causes the same responses as perceiving the corresponding stimulus (for review, see Hesslow, 2002). The second concept is top-down regulation of sensory input (Frank, 2016; Lopresti-Goodman et al., 2013), suggesting if mental representations of stimuli are generated and cognitively manipulated, the perception of such stimuli is subjected to top-down adaptation. In other words, forming mental representations of stimuli, between perceiving and responding to them, allows for top-down downregulation of sensory input. For example, if participants are asked to judge whether they need one or two hands for lifting planks of different sizes while they either grasp them or only look at them, the latter condition but not the former will be subjected to top-down sensory adaptation (Lopresti-Goodman et al., 2013). The third concept is predictive coding (Friston, 2010), which suggests that any action, that is, motion or perception, will be initiated by forming predictions about the next state of the motor and/or sensory apparatus (Adams et al., 2013; Clark, 2013). Next, predictions are propagated downward through cortico-cortical and corticospinal projections to the relevant muscles and sensory units. Notably, down propagating signals are always *predictions* and not *motor commands* (for the treatment of neuromuscular mechanisms, see Adams et al., 2013). If there will be a difference between the state of the system and the prediction, a prediction error is formed (i.e., being in the surprise state). Any self-organizing system aims to reside in the lowest possible energy state. Therefore, in the surprised-state, such systems attempt to minimize prediction errors (Friston, 2010). During volitional movements, predictions have a higher weight in comparison to prediction errors. Consequently, to leave the surprise state, prediction errors are used in reflex arcs to correct the movement and align it with predictions. Hence, prediction errors are gradually downregulated in reflex arcs during backpropagation and are diminished sufficiently not to be propagated beyond thalamic nuclei (Adams et al., 2013; Brown et al., 2013). During perception, on the other hand, prediction errors are given a higher weight compared to predictions. Hence, this time, predictions are updated based on prediction errors for coming out of the surprised-state. In predictive coding theory, these two processes are called active and perceptual inference, respectively. Despite its popularity and success in explaining normal perception and action, predictive coding cannot explain, why during hypnosis, participants can execute actions described by suggestions but attribute the action to external sources, that is, perceive them to be caused externally rather than by their own volition (Lynn et al., 1990). In other terms, in terms of the predictive coding model, to start a movement, predictions must be given a higher weight than sensory feedback from the external world (i.e., active inference); however, participants will only perceive their response as externally originated if prediction errors have a higher weight than predictions (i.e., perceptual inference) (Brown et al., 2013; Clark, 2013). There are several different theoretical attempts to explain the sense of involuntariness during hypnotic-induced actions via the predictive coding model (e.g., Jamieson, 2016; Martin & Pacherie, 2019); however, here, we will only focus on one presented by SATH.

SATH employs three top-down cognitive processes to explain hypnotic and posthypnotic suggestion-induced responses. (1) *Cognitive simulation* (Farah, 1988; Hesslow, 2002). During hypnosis, participants have two sources of input, cognitive simulations and the external world. In other words, besides perceptual input from the external world, stimuli described by suggestions that are cognitively simulated by participants provide “perceptual” input. (2) *Sensory adaptation* (Frank, 2016; Lopresti-Goodman et al., 2013). When sensory input from external stimuli is not aligned with cognitive simulations and suggestions, top-down sensory adaption may downregulate perceptual input from the external world. Together with predictive coding, cognitive simulation and sensory adaptation can offer an explanation of how hypnotized participants execute responses and attribute them to external sources. If one assumes that during hypnosis, cognitive simulations are given higher weight in comparison to predictions, then predictions can be updated based on perceptual input from cognitive simulations (perceptual inference). Simultaneously, predictions are given a higher weight in comparison to sensory input from the external world. Therefore, even though sensory input from the external world is used in reflex arcs to align movements with predictions (active inference), it is simultaneously downregulated by top-down sensory adaptation, preventing it from passing beyond thalamic nuclei. Consequently, hypnotized participants exert a suggested motion and simultaneously attribute it to the suggestions (or external sources) rather than themselves. (3) *Mental practice and problem-solving* (Zahedi, Sturmer, et al., 2020). There are two situations that cannot be explained by cognitive simulation and/or top-down sensory adaption and, as a result, require a further cognitive top-down process. (I) When suggestions do not provide cognitive-simulation-provoking descriptions of stimuli, hypnotized participants need to fill in the gap and find an appropriate strategy to simulate presented stimuli cognitively. (II) In case that suggestions aim to form a new trigger-response contingency, cognitive simulations can provide an (imagined) exercise environment, where new strategies, either outlined by suggestions or developed by participants themselves, are mentally practiced.

Notably, SATH hypothesizes that the above-mentioned cognitive processes can be used to different extents and in various combinations, depending on the individual’s capabilities of participants. For instance, some participants might heavily rely on cognitive simulation and others on top-down sensory adaptation. This proposition is in line with findings, showing there are different groups of the high-hypnotic-suggestibles who might rely distinguishably on different cognitive processes for responding to suggestions (Terhune, 2015; Terhune & Hedman, 2017; Terhune & Oakley, 2020).

According to SATH, besides the aforementioned top-down cognitive processes, social and psychological factors may also be of great importance. Since all the top-down processes above are volitional and goal-directed, participants’ expectations, openness, and willingness will determine whether they will be motivated to engage in the responses described in the suggestions. It is crucial to notice that the presented cognitive processes are hypothesized by SATH, and even though based on previous studies and results, they are theoretically justified (c.f., Zahedi & Sommer, 2021), their relevance should be corroborated by novel empirical investigations.

### 1.1. Theory-driven Hypotheses for CFA and SEM

In the present study, we used SATH as a framework to formulate theory-driven hypotheses to be tested by CFA and SEM. Based on SATH, a bifactor model should best explain the variance in the HGSHS:A scores. The proposed bifactor model consists of a G-factor and three minor factors, as will be justified next and illustrated in Figure 1.

**Figure 1.**
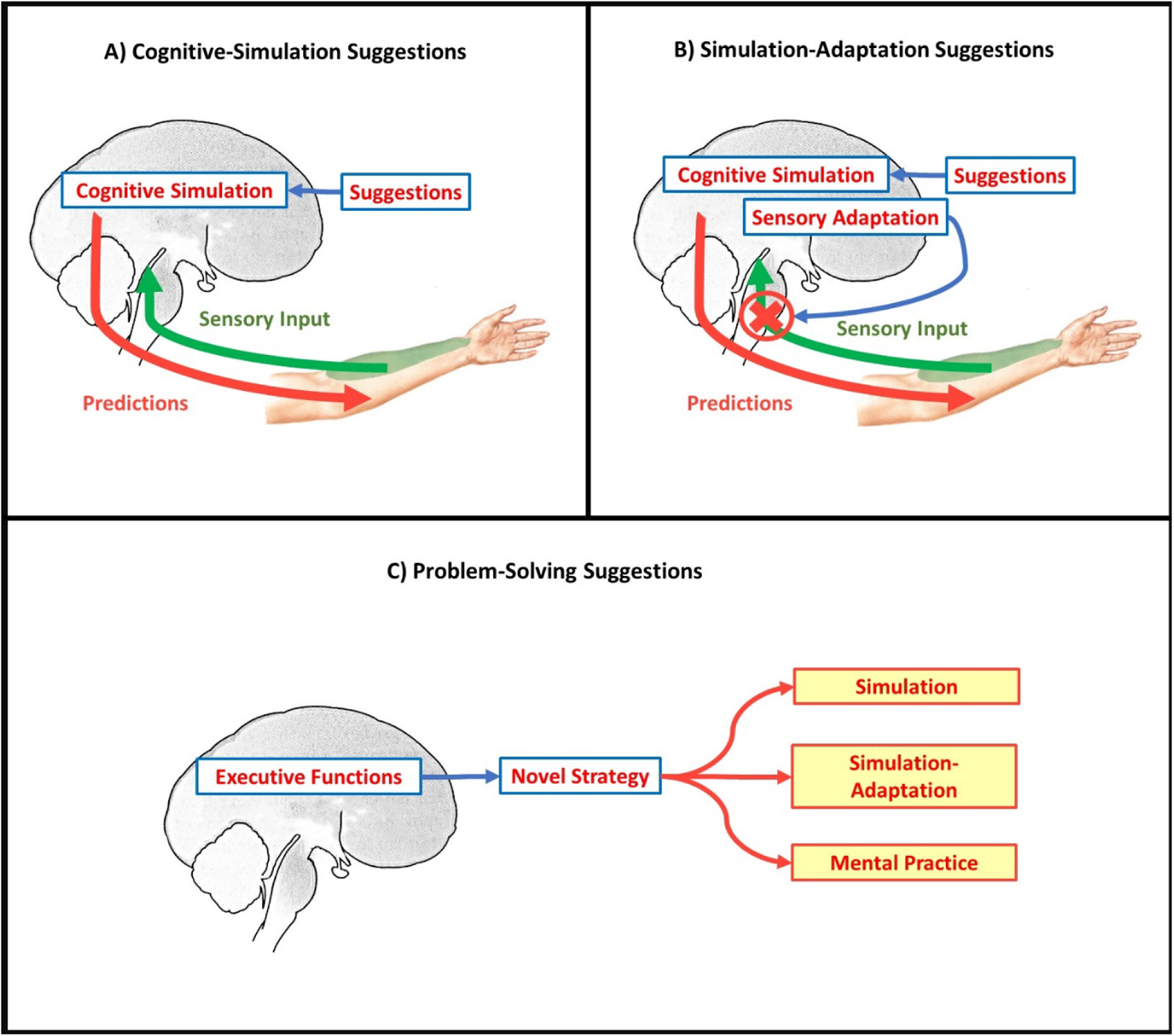
Schematic representation of different kinds of hypnotic and posthypnotic suggestions and the hypothetical underlying processes; A) cognitive-simulation, B) simulation-adaptation, and C) problem-solving suggestions. For details, please see text.

The first minor factor (direct-ideomotor) covers cognitive-simulation suggestions, where sensory information is congruent with the portrayed response (Fig. 1A). For instance, consider the suggestion: “stretch your arm and keep it in the air, after a while your hand starts to feel fatigued and it starts to move downward. It is as if a heavyweight has been put on your arm” (Shor & Orne, 1962). Here, sensory input is aligned with the suggestion; if participants stretch their arms, they will feel fatigued after a while. Therefore, the suggestion predicts a sensation that will indeed occur. Nonetheless, if the suggestion is successful, the fatigue will not be attributed to body-internal processes (Kihlstrom, 2008; Lynn et al., 1990) but to the hypnotic condition. Based on SATH, this will only happen if participants cognitively simulate what is described by direct-ideomotor suggestions (i.e., a heavyweight is put on their arm). Hence, cognitive simulation is necessary for the successful exertion of targeted hypnotic-suggestion-induced responses.

The second group of suggestions (challenge-ideomotor) can be described as simulation-adaptation suggestions (Fig. 1B). In these suggestions, sensory information is incongruent or conflicting with the suggested information. For instance, consider the hand levitation suggestion: your hand feels lighter “as if there’s a large helium balloon under [your] palm, or attached with strings to each [one of your] fingertip and [your] wrist … [your] hand and arm will begin to float up” (Hammond, 1998, pp. 43-44). As explained by SATH, when sensory input is not aligned with cognitive simulation, it is downregulated through the top-down mechanism of sensory adaptation. Together, cognitive simulation and sensory adaptation can explain why the motion described in the suggestion is executed by the hypnotized participant, but nevertheless, attributed to an external cause. Let’s consider the hand levitation suggestion. Here, predictions are generated based on cognitive simulations of the suggestions’ description (i.e., helium balloons are attached to the fingers). Since the predictions (i.e., the hand and arm will be levitated) will receive a lower weight than the cognitively simulated input, they are updated based on prediction errors formed from the comparison of predictions with cognitive simulation feedback. This explains why the motion is attributed to an external source (i.e., helium balloons). At the same time, predictions will be given a higher weight compared to sensory feedback, which is downregulated by top-down sensory adaptation. Consequently, the targeted motion is executed, and sensory feedback will be used for adapting the movement with predictions. Evidently, in challenge-ideomotor suggestions, besides cognitive simulation, sensory adaptation must be incorporated. This fact distinguishes this category from direct-ideomotor suggestions, where only cognitive simulation is necessary.

The third minor factor (perceptual-cognitive) relates to suggestions that require problem-solving, for example, in order to find an appropriate strategy for cognitive simulation, forming a new trigger-response contingency, or adapting an existing response to a novel situation. In the most common cases, a problem-solving suggestion will only describe a goal but no concrete strategy for accomplishing the goal, and participants are responsible for filling the gap. That is, participants have to find an appropriate cognitive strategy and then implement the necessary mechanisms for executing this strategy. Importantly, after finding a suitable strategy for responding to a suggestion, the suggestion will turn into a direct-ideomotor or challenge-ideomotor suggestion (Fig. 1C). Consider, for example, the posthypnotic suggestion item of HGSHS:A: after the termination of hypnosis, “when you hear a tapping noise, you will reach down and touch your ankle” (Shor & Orne, 1962). The suggestion has clearly defined the goal but no strategy for implementing it. If participants merely reach down and touch their ankle while thinking that they are doing this because the suggestion instructed or commanded them to do so, the action will be attributed to the exertion of direct volitional effort. However, in an alternative scenario, if participants imagine the sound and repeatedly connect it to itching or burning in their ankle, they will exert the portrayed action after hearing the sound but will not attribute it to any volitional effort of their own. Any suggestion can be a perceptual-cognitive suggestion if it fails to imply an applicable strategy for the suggested action. The notion that the content of suggestions, and not only the action described by them, can also affect participants’ responses has been confirmed by Galea et al. (2010). In their study, first, high-hypnotic-suggestible participants were given a suggestion, aiming to induce rigidity and stiffness in their arms. Afterward, participates were asked to move their arms with the implication that they cannot do it. The authors intentionally did not present a relevant strategy (for instance, believing to be paralyzed). As inferred by electromyography, the participants of Galea et al. (2010) came up with different strategies, such as simultaneously activating agonist and antagonist muscles (biceps and triceps), only activating the antagonist (triceps) but inactivating the agonist, or not activating any muscle.

Based on SATH (c.f., Zahedi & Sommer, 2021), we assume that simple multifactor models are not adequate for explaining the variance in hypnotic-susceptibility scales. Instead, two alternatives to the multifactor model presented in Table 1 can be proposed. First, in contrast to other top-down cognitive processes required for responding to suggestions, SATH assumes that cognitive simulation is needed to successfully execute all types of suggestions presented in HGSHS:A. This proposition is corroborated by the meta-analysis of Landry et al. (2017), which showed that the activation of areas in brain commonly related to imagination, such as the lingual gyrus (Jung et al., 2016), is the shared characteristic of many different forms of suggestions. Hence, one might expect, treating the direct-ideomotor factor (i.e., a minor factor) as a G-factor should improve the multifactor model. In other words, this model tests whether one can assume cognitive simulation as the main cognitive process needed for all items of HGSHS:A and other cognitive processes as secondary and additive.

The second model focuses on SATH’s hypothesis that there are two sources of variability in hypnotic suggestibility, echoing the proposition of Kirsch (1997), asserting that hypnotic suggestibility can be decomposed into general suggestibility (capability of responding to suggestions regardless of hypnosis) and hypnotizability (increase in suggestibility due to hypnotic induction). Notably, by general suggestibility, we are referring to direct verbal suggestibility, which should be distinguished from other forms of suggestibility (Oakley et al., 2021; Polczyk, 2016). Hence, besides the specific correlated minor factors, capturing the different suggestibilities described above (related to different top-down cognitive processes), one may expect that the addition of a general factor to which all HGSHS:A items contribute will improve the model. Evidently, two G-factors are included in this model. That is, the suggestibility G-factor, measured by the correlated minor factors, and the G-factor of hypnotizability, capturing the psychosocial variables involved in responding to hypnotic suggestions (i.e., hypnotizability). Notably, the G-factor of hypnotizability is unrelated to cognitive processes involved in responding to suggestions (i.e., the suggestibilities). In other words, in this model, four latent factors are included, that is, three minor factors and one G-factor. It is observable that both Kirsch (1997) and SATH have a hidden assumption; the same cognitive processes that are employed for responding to suggestions outside of hypnosis will also be employed under the influence of hypnosis. Therefore, any alteration in suggestibility is related to psychosocial variables (which can be different inside and outside of hypnosis).

Finally, an important question regarding HGSHS:A is whether the outcome of one category of suggestions predicts the outcomes of other categories. Based on SATH, the challenge-ideomotor factor encompasses items that require a combination of the essential underlying mechanisms (i.e., cognitive simulation and sensory adaptation) for complying with suggestions. Consequently, the outcome of the challenge-ideomotor factor might predict success in the direct-ideomotor factor and, to a lesser extent, also in perceptual-cognitive suggestions. In other words, we expect the challenge-ideomotor outcomes to predict the direct-ideomotor and cognitive-perceptual results, but not vice versa.

Together, in the current study, for solving several issues related to EFA, we used CFA and SEM to investigate the structural construct of the HGSHS:A scores. Since CFA needs a theory from which relevant hypotheses can be derived, after scrutinizing possible candidates, we chose the SATH and used it as the basis for our CFA and SEM. Based on this theory, a bifactor model was postulated, consisting of three minor factors, each measured by a distinguished group of suggestions, and a G-factor, measured by all items. Further, the internal construct of minor factors was investigated to establish a predictive pathway between categories of suggestions.

## 2. Methods

### 2.1. Participants

A sample of 477 participants ( 252 *Women, manage* = 28.7 *years*, *SD* = 12.6 *years*) was recruited. Several different methods have been used for finding prospective participants; besides inviting local Psychology students, the study was advertised on eBay Kleinanzeigen (https://www.ebay-kleinanzeigen.de) and local radio stations. The study had been approved by the ethics committee of the Humboldt-Universität zu Berlin. Prior to the experiment, signed consents were obtained. Participation was compensated by free assessment of hypnotic susceptibility or course credits.

Based on the most conservative estimation, the sample size of a CFA should be *N* = 20 * *free parameters* (Tanaka, 1987). In our study, the number of free parameters in the basic model was 24, which shows our sample was big enough, *N* ≈ 20 * 24. Further, based on 1- β > 0.9, *H*0 < 0.05, and *H*1 > 0.1 (Cohen, 2016), we calculated the required sample size for Root Mean Square Error of Approximation (RMSEA) (Preacher & Coffman, 2006), which yielded a minimum required sample size of, *N* = 120 which again confirms that the analyses conducted here have sufficient power to test presented hypotheses.

### 2.2. Measurements and Procedure

HGSHS:A (Shor & Orne, 1962) has 12 suggestions (Table 1) and is designed for administration in group sessions. In the present study, we screened groups of 2-15 volunteers per session. At the beginning of the session, a short description of hypnosis and hypnotizability was given by a certified hypnotizer (A.Z.), as advised in the HGSHS:A manual (Shor & Orne, 1962). Afterward, a recorded German version of HGSHS:A (Bongartz, 1985) was administered while participants were required to follow the suggestions. After the presentation of all suggestions, participants completed a questionnaire regarding their experience, consisting of two sections. (1) An objective section inquired whether the participant had complied with each of the suggestions, and (2) a subjective section asked how strongly they had experienced the effect of each suggestion. For recording the objective and subjective responses, the German booklet of HGSHS:A was used (Bongartz, 1985). Notably, even though called objective, the objective items of HGSHS:A are still responded to by participants and are examples of self-report. In the objective section, responsiveness or non-responsiveness was scored as 1 or 0, respectively, for the first 11 suggestions; the 12^th^ suggestion (i.e., posthypnotic amnesia) was scored as 1, if less than four items could be remembered in the initial amnesia test and more than one item in the reversibility test, and otherwise as 0. This scoring is based on the suggestions of Kihlstrom and Register (1984), which is used widely in the field. Hence, participants could receive a total score of 0 =< *hypnotic suggestibility* =< 12.

Noticeably, in the objective section, for each suggestion, an observable criterion is presented on which participants should base their responses. For instance, “[w]ould you estimate that an onlooker would have observed that your hand lowered at least six inches (before the time you were told to let your hand down deliberately)?” Participants’ responses are independently verified by the hypnotist and administrators present at the session. In case that there were significant deviations between participants’ responses and administrators’ observations, the participant’s data were discarded. However, less than 5 participants’ responses were discarded according to these criteria.

### 2.3. Data analyses

All data analyses were conducted in R (R Core Team, 2013); for CFA and SEM, the lavaan package was used (Rosseel, 2012). As the objective scores of HGSHS:A are binary, the diagonally weighted least squares (DWLS) was used to estimate model parameters, and the full weight matrix (WLSMV) was utilized to compute robust standard errors and mean- and variance-adjusted test statistics. Since the study benefitted from a large sample, WLSMV was preferred for ordinal data in comparison to maximum likelihood (ML) or robust maximum likelihood (MLR) (Li, 2016). Distributions were fitted with VGAM (Yee, 2010; Yee, 2015) and fitdistrplus (Delignette-Muller & Dutang, 2015). In order to evaluate whether the variance in the HGSHS:A scores can be attributed merely to the variance in the difficulties of items, the Rasch model was applied before conducting CFA. Rasch models were calculated by eRM (Mair & Hatzinger, 2007) employing conditional maximum likelihood (CML) and tested using Andersen’s likelihood-ratio test (Andersen, 1973).

## 3. Results and Discussion

### 3.1. Descriptive results

The Cronbach’s alpha coefficients and 95% confidence intervals for the objective ( *mean* = 6.37[6.15 - 6.60]; *SD* = 2.49 ) and subjective items ( *mean* = 61.68[59.57 - 63.79]; *SD* = 22.84 ) were acceptable, *Cronbach’s α_Objective items_* = .67 [.61 - .69] and *Cronbach’s α_Subjective items_* = .90 [.88 - .91], respectively.

Figure 2 presents the distribution of the HGSHS:A scores; since these parameters are ordinal, they cannot be expected to have a normal distribution. However, the methods used for both CFA and SEM analyses and calculation of estimated loadings in CFA and SEM are robust and insensitive to deviations from normality. Regarding the distribution of the HGSHS:A scores, three points should be noted. First, the scores show a beta-binomial distribution with estimated μ = 0.52 and ρ = 0.099^2^ (the distribution fit indices are presented in Table 5). Second, it could be argued, as in our study, participants were volunteers, the sampling procedure might have been biased toward more hypnotic-susceptible participants. However, the distribution is not biased (skewed) toward the low- or high-hypnotic-suggestibles. Third, the bimodal shape of the distribution is of great interest and will be discussed in section 3.3.

**Figure 2.**
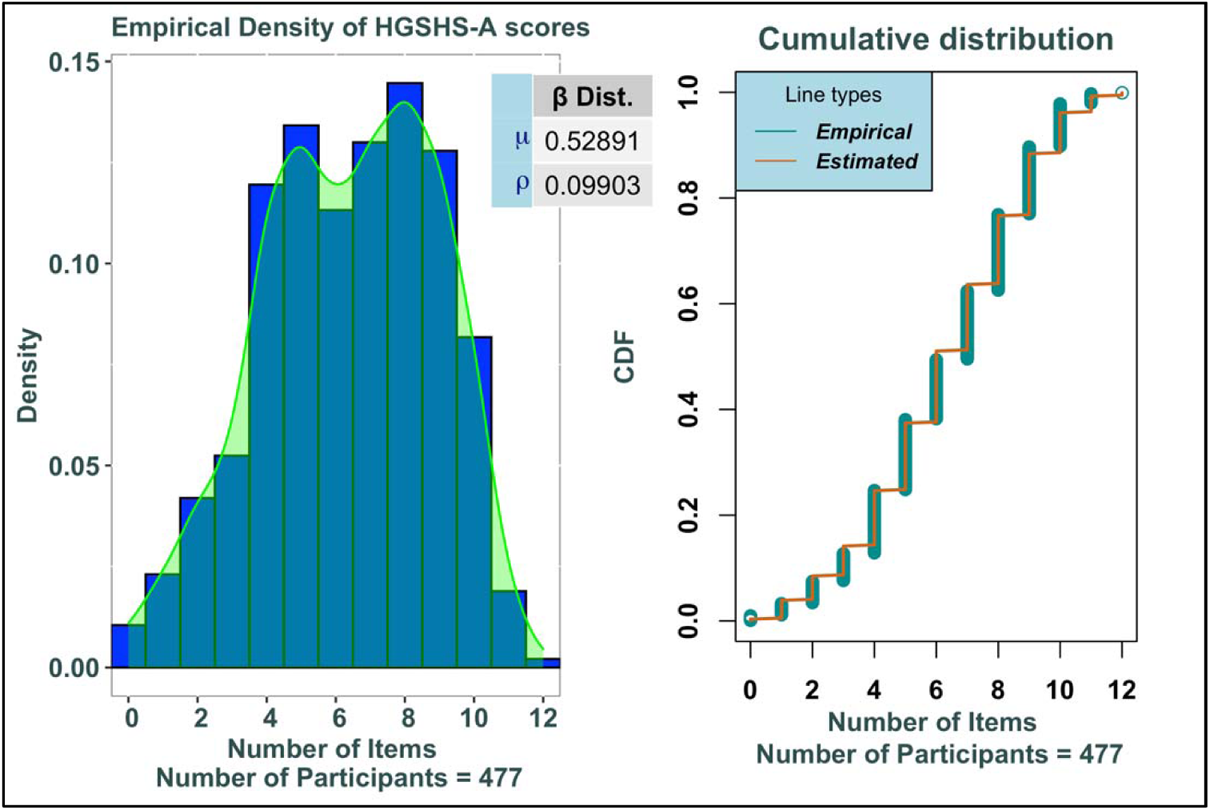
Left: Histogram and density estimation distribution of the HGSHS:A scores. Right: Cumulative distribution of the HGSHS:A scores.

For testing the global fit of the Rasch model, scores were split based on median, and Andersen’s likelihood-ratio test results showed bad fit (χ^2^(11) = 70.802, b < 0.001)^3^. This result indicates that the heterogeneity in HGSHS:A cannot be attributed to variance in item difficulties alone. This conclusion is in agreement with the findings of other Rasch analyses, such as McConkey et al. (1980), investigating HGSHS:A, and Näring et al. (2004), assessing SHSS:C. In other words, these scales cannot be considered unidimensional. Notably, the item characteristic curves (ICC) (Fig. 3A) and the wright map of HGSHS:A items (Fig. 3B) showed that items related to the direct-ideomotor (*mean* β = −1.161), challenge-ideomotor (*mean* β = − −0.049), and perceptual-cognitive factor (*mean* β = 1.629) differ in their difficulty.

**Figure 3.**
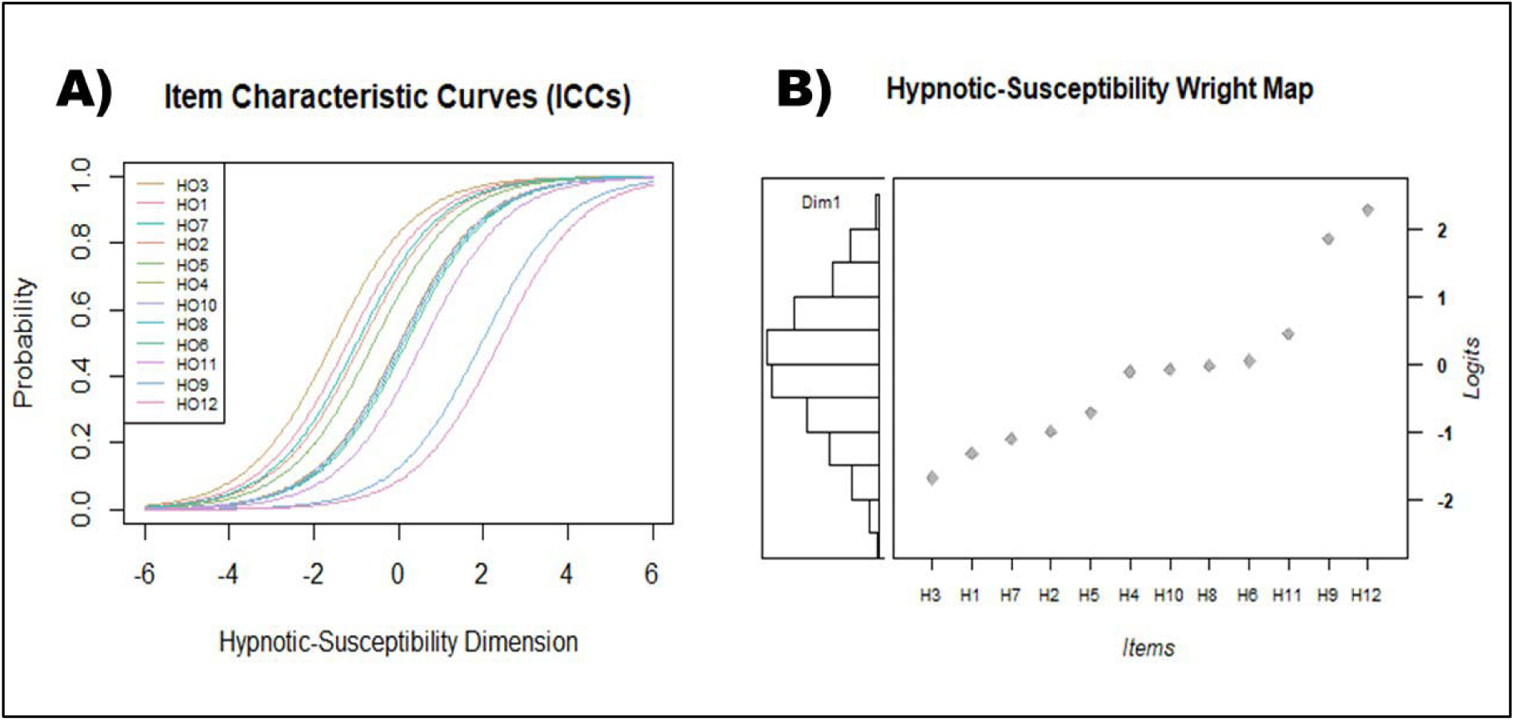
The results of Rasch analysis of HGSHS:A scores. A) The item characteristic curves (ICC) for each of the items; B) The item-person diagram (i.e., wright map) for all items (right) and participants (left).

### 3.2. Confirmatory Factor Analyses

For testing the hypotheses outlined in section 1.1, we fitted four models to the data (Fig. 4) (several other modes were also tested. For details, see the supplementary material). First, we tested a single-factor model, with the hypothesis that there might be a single G-factor, which can account for all the variance in the data. Model 2 (i.e., the three-factor model) represents the basic multifactor model introduced in Table 1, where each suggestion is related to only one of the three latent factors.

**Figure 4.**
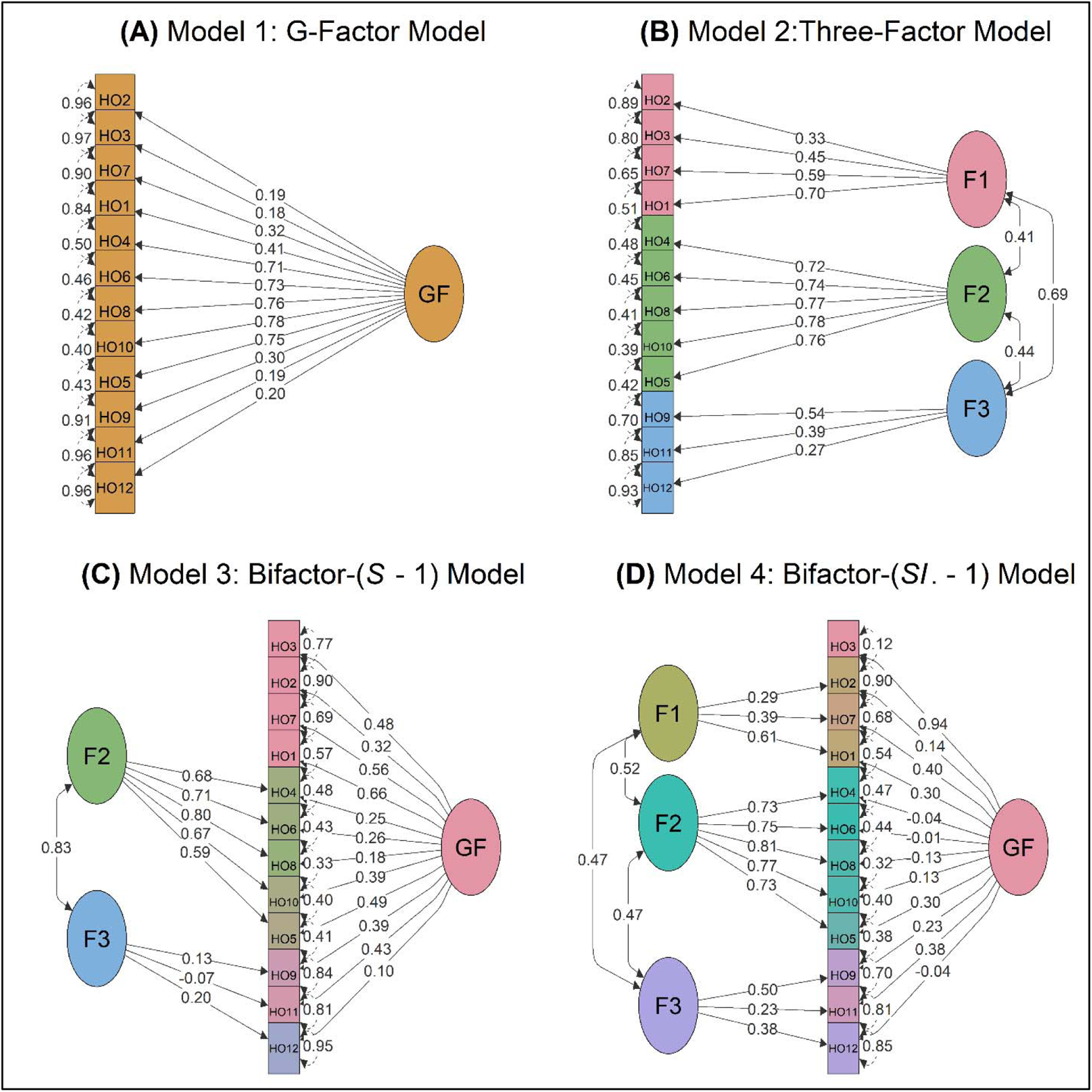
Estimated loadings of the CFA models. (A) G-factor model (#1), (B) three-factor model (#2), (C) bifactor- (*S* − 1) model (#3), and (D) bifactor- (*S.I* − 1) model (#4). On the single-headed arrows, standardized factor loadings are given. All loadings in the G-factor and three-factor models are significant, *p* < .05. In the bifactor- (*S* − 1) model (Model 3), the loadings of Factor 3 and HO12 on GF are not significant; however, other loadings are significant, *p* < .05. In the bifactor- (*S.I* − 1) model (Model 4), the loadings of HO2, HO4, HO6, HO8, HO10, and HO12 on the G-factor are not significant; however, other loadings are significant, *p* < .05. Loadings are equivalent to standardized regression coefficients (beta weights), and they are estimated with diagonally weighted least squares. The self-loops show error terms. Squaring these terms gives an estimate of the variance for each item that is not accounted for by the latent construct. The curved, double-headed arrows indicating correlation coefficients between latent variables, all of which, except the correlation between F3 and F1 in bifactor- (*S* − 1) model (Model 3), are significant, *p* < .05. *Note:* F1: direct-ideomotor suggestions; F2: challenge-ideomotor suggestions; F3: perceptual-cognitive suggestions.

Models 3 and 4 represent two bifactor models that were proposed in section 1.1. to enhance the basic multifactor model. In conventional bifactor models, it is necessary to assume orthogonality between latent factors. That is, correlations between latent factors are constrained to zero (Reise, 2012). Otherwise, interpretation may be complicated (Musek, 2017; Reise, 2012), and models may be unidentifiable (i.e., common anomalies of bifactor models) (Eid et al., 2017; Eid et al., 2018). Anomalies in conventional bifactor models may arise if data were not obtained in a two-level sampling procedure (Eid et al., 2017); that is, if not both, participants, as well as items to participants, have been randomly assigned. Such a two-level sampling is not possible for HGSHS:A since items are fixed. Therefore, based on the suggestions of Eid et al. (2017), we computed bifactor- (*S* − 1) and bifactor- (*S.I* − 1) models instead of conventional bifactor models. In the bifactor- (*S* − 1) model (Model 3), one of the minor factors (i.e., direct-ideomotor) was conceptualized as a G-factor measured by all HGSHS:A items. The reason for choosing direct-ideomotor as the reference domain was that, based on SATH, cognitive simulation is a top-down process required for responding to different forms of suggestions. Sensory adaptation and problem-solving, in contrast, are necessary only for a special group of suggestions. In the bifactor- (*S.I* − 1) model (Model 4), in addition to three *correlated minor factors*, a G-factor measured by all HGSHS:A items was assumed. In this model, one item is reserved only for the G-factor. This indicator serves to distinguish the G-factor from the minor factors. The reason for choosing Item 3 of HGSHS:A as the scaling indicator for the G-factor is twofold. First. This item is responded to by most participants (Bongartz, 1985; Shor & Orne, 1963). Therefore, despite depending on hypothesized psychosocial variables like every other item of HGSHS:A, it should require more accessible cognitive processes compared to other items. Second, Item 3 is the first item of HGSHS:A that is presented after the hypnotic induction. Consequently, Item 3 is an appropriate scaling indicator for a G-factor, which assumingly taps into psychosocial variables. The choice of Item 3 as the scaling item was a priori decision.

Table 2 summarizes the fit indices of all four models, graphically presented in Figure 4. Further, the reliability coefficients of factors in all models are presented in Table 3. Based on the results shown in Table 2, and considering adjusted thresholds, the G-factor model (Model 1) showed a poor fit to the data. The three-factor (# 2) and bifactor- (*S* − 1) (# 3) models fared better and had a lower standardized root-mean-squared residual (RMSEA) compared to Model 1; however, they still showed only a modest fit. In contrast, the bifactor- (*S.I* − 1) (# 4) model showed a perfect fit to the data. The bifactor- (*S.I* − 1) model has the lowest χ^2^ and standardized root mean squared residual (SRMR), indicating a better fit than other models, which is expected, as more complex models usually fit the data better.

**Table 2.** Fit Indices of the models assessed by confirmatory factor analysis (CFA)

| Model | *df* | $\chi^2$<sup>a</sup> | RMSEA [90% CI]<sup>b</sup> | SRMR<sub>c</sub> | CFI<sup>d</sup> | TLI<sup>d</sup> |
| --- | --- | --- | --- | --- | --- | --- |
| Model 1: G-factor model | 54 | 169.93 *** | 0.067 [0.056-0.079] | 0.105 | 0.887 | 0.861 |
| Model 2: Three-factor model | 51 | 99.06 *** | 0.045 [0.031-0.058] | 0.079 | 0.953 | 0.939 |
| Model 3: Bifactor- (*S* – 1) model | 45 | 80.95 ** | 0.041 [0.026-0.055] | 0.068 | 0.965 | 0.948 |
| **Model 4: Bifactor- (*S*, *I* – 1) model** | **40** | **50.7** | **0.024 [0.000-0.042]** | **0.048** | **0.99** | **0.983** |

*Note:* The endorsed model is indicated in bold. SRMR: standardized root-mean-squared residual; RMSEA: Root Mean Square Error of Approximation; CFI: Bentler's Comparative Fit Index; TLI: Tucker-Lewis Index.
<sup>a</sup> When $\chi^2$ test is not significant the model fits the data. However, as the $N = 477$ was very large, it is expected that $H_0$ would be over-rejected.
<sup>b</sup> Lower values of RMSEA indicate better fit, with values $< .05$ indicating a close fit to the data (Xia & Yang, 2019). For $1 - \beta > 0.9$ , $H_0 < 0.05$ , and $H_1 > 0.1$ the required sample size is $N > 120$ .
<sup>c</sup> Lower values of SRMR indicate better fit, with $\text{SRMR} < .08$ indicating a close fit to the data.
<sup>d</sup> Values $> .95$ for CFI and TLI indicate a good fit (Xia & Yang, 2019). TLI is not normalized and may have values $> 1$ .
\* $p < .05$ ; \*\* $p < .01$ ; \*\*\* $p < .001$

**Table 3.** Ordinal coefficient alpha *^a^* of latent factors of all models.

| Model | F1 | F2 | F3 | G-Factor |
| --- | --- | --- | --- | --- |
| Model 1: G-factor model | - | - | - | 0.76 |
| Model 1: Three-factor model | 0.58 | 0.86 | 0.35 | - |
| Model 1: Bifactor- ($S - 1$) model | - | 0.86 | 0.35 | 0.76 |
| Model 1: Bifactor- ($S, I - 1$) model | 0.51 | 0.86 | 0.35 | 0.76 |

*Note:* for the details of the models, see the text.
F1: direct-ideomotor suggestions; F2: challenge-ideomotor suggestions; F3: perceptual-cognitive suggestions.
<sup>a</sup> As hypnotic-susceptibility scores are dichotomous, the ordinal coefficient alpha (Zumbo et al., 2007; Zumbo & Kroc, 2019) was calculated.

Even though only the bifactor- (*S.I* − 1) model (# 4) closely fits the data, we compared all four models using the likelihood ratio test (Tab. 3). We reasoned that, since we had a large number of participants, χ^2^ might over-reject null hypotheses (*H0*: a given model does not fit the data). All of our models can be tested using the likelihood ratio test, as suggested by Reise (2012). The likelihood ratio tests were conducted hierarchically; that is, a model was only compared to the next simpler model. The results of these tests (Table 4) confirm that the G-factor model (Model 1) is least capable of capturing the variance in the data. Hence, HGSHS:A is definitely not a homogenous scale as it might be assumed, based on the fact that no subscale is defined for it. This becomes clear if one notices that most commonly, the general score of HGSHS:A is used for predicting participants responsiveness to suggestions outside of screening sessions (e.g., Zahedi et al., 2019; Zahedi, Luczak, et al., 2020; Zahedi et al., 2017; Zahedi, Sturmer, et al., 2020). Second, the bifactor- (*S* − 1) model (# 3) fits the data significantly better than the three-factor model (# 2), corroborating our theoretically derived hypothesis that cognitive simulation is the shared top-down cognitive process employed in all forms of suggestions. Finally, the bifactor- (*S.I* − 1) model (# 4) is significantly better in comparison to Model 3, supporting our hypothesis that two sources of variability are affecting hypnotic-susceptibility scores simultaneously (which, based on our hypotheses, can be related to hypnotizability and suggestibility).

**Table 4.** Results of likelihood ratio tests comparing all models

| Competing models | | $df$ | $\chi^2{}^a$ |
| --- | --- | --- | --- |
| Model 2: Three-factor model | Model 1: G-factor model | 3 | 47.9 *** |
| Model 3: Bifactor- ($S - 1$) | Model 2: Three-factor model | 6 | 18.9 ** |
| Model 4: Bifactor- ($S.I - 1$) | Model 3: Bifactor- ($S - 1$) | 5 | 22.2 *** |

*Note:* <sup>a</sup> If the $\chi^2$ test is significant the more complex model will be endorsed, if not, the simpler model is endorsed.
\*\* $p < .01$ ; \*\*\* $p < .001$

With regard to Model 4, one should consider that a model with three correlated first-order factors without a G-factor is equivalent to a second-order G-factor model; that is, a model in which, besides three minor uncorrelated factors, a second-order G-factor is assumed (Eid et al., 2017). Hence, in our bifactor- (*S.I* − 1) model (# 4), two G-factors, one hidden and one explicit, are assumed, which capture variance from two different sources. In the introduction, we argued that based on a proposal by Kirsch (1997), which is also incorporated in SATH (Zahedi & Sommer, 2021), hypnotic suggestibility, as measured by HGSHS:A, can be decomposed into suggestibilities (capability of responding to different types of suggestions in general) and hypnotizability (the alteration in suggestibilities due to the hypnotic condition). Consequently, in our bifactor- (*S.I* − 1) model (# 4), the correlated minor factors measure suggestibilities, and the G-factor measures hypnotizability. Notably, based on the presented hypotheses, we did not assume orthogonality between minor factors themselves, as we expected that cognitive processes would be correlated. Hence, in bifactor models, only the correlations between the G-factor and minor factors were restricted to zero. Although suggestibility and its components can be clearly defined by SATH, hypnotizability is less identifiable and can be related to various psychosocial variables, including willingness and openness (Green & Lynn, 2011; Lynn et al., 2015), the prior expectations about hypnosis (Kirsch & Lynn, 1997; Terhune et al., 2017), expectations induced by the wordings of suggestions (Lynn et al., 1987; Matthews et al., 1985; Spanos, 1971), rapport with the hypnotist (Lynn et al., 2019), and motivation to respond to suggestions (Jones & Spanos, 1982).

Two critical points related to the loadings in the bifactor- (*S* − 1) and (*S.I* − 1) models should be discussed here. Firstly, in Model 3, problem-solving indicators loaded significantly on the G-factor but not on the problem-solving factor. This result can be interpreted post hoc. That is, as in section 1.1. we argued that for perceptual-cognitive suggestions, participants need first to find an appropriate responding strategy, and then these suggestions turn into typical direct-ideomotor or challenge-ideomotor suggestions. As each of these suggestions poses a unique problem, it seems that cognitive simulation (the G-factor) is more salient than problem-solving in responding to these suggestions. Secondly, in Model 4, simulation-adaptation indicators loaded strongly on the direct-ideomotor factor but very weakly on the G-factor. One interpretation of this finding is that as the challenge-ideomotor suggestions require two essential top-down cognitive processes, motivations, expectations, and other psychosocial variables are less relevant or irrelevant in responding to them. However, one can also explain the observed results by assuming a cognitive mechanism related to some specific motor-related suggestions, which is not shared by other suggestions. It should be noted that these anomalies in the bifactor models were not expected, and therefore, the presented interpretations regarding these anomalies should be assessed empirically in future studies. Further, even though Item 2 loads significantly on predicted factors, the loadings are not strong. This, however, can be related to the context and nature of the item since, in other studies, similar issues were encountered during EFA (for review, see Sadler & Woody, 2021).

### 3.3. Structural Equation Modeling

Next, we tested our hypotheses that the specific minor factor, which is parceling the suggestions with all the essential top-down processes required for different suggestibilities, can predict the outcome of the other minor factors. In other words, we expected that the challenge-ideomotor factor can predict the direct-ideomotor and cognitive-perceptual factors, but not vice versa. We tested this hypothesis in the models with three minor factors (Models 2 and 4; Fig. 5). Both the direct-ideomotor and cognitive-perceptual factors regressed significantly to the challenge-ideomotor factor, bi < .05. However, in none of these models, the challenge-ideomotor factor regressed significantly to the direct-ideomotor or cognitive-perceptual factors, bi > .1 model), the correlation between the direct-ideomotor and cognitive-perceptual factors became non-significant when they regressed on the challenge-ideomotor factor, indicating that relationships between these factors can be reduced to regression of the perceptual-cognitive (problem-solving suggestions) and direct-ideomotor (cognitive-simulation suggestions) on the challenge-ideomotor factor (simulation-adaption suggestions).

**Figure 5.**
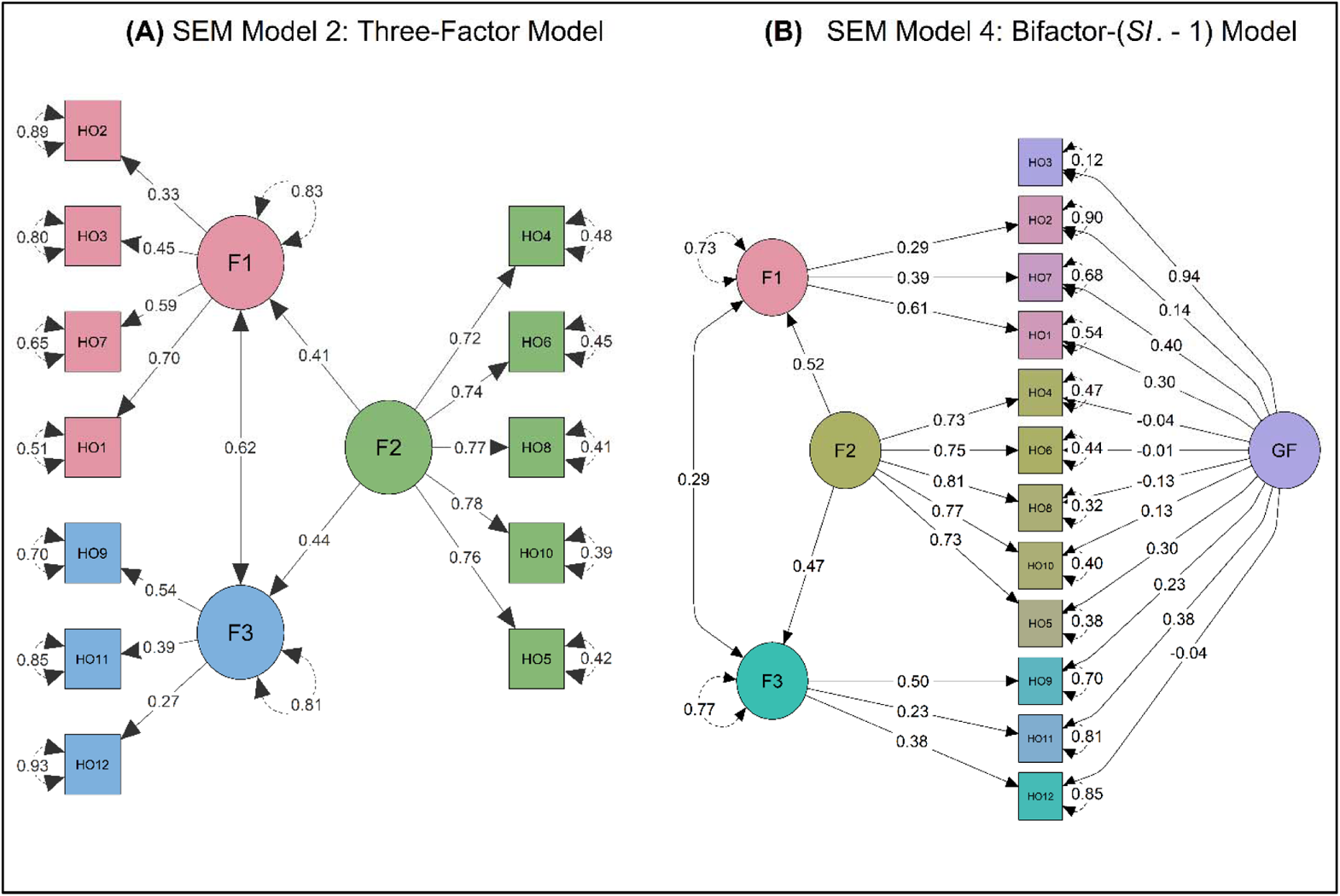
The structural segment of (A) SEM Model 2; the three-factor, and (B) SEM Model 4: bifactor- (*SI*. − 1) models. All correlation and regression coefficients between latent factors, except the correlation between t.3 ~~ t.1 in the bifactor- (*SI*. − 1) model, are significant, *p* < .05. For statistical details of loadings of items on latent factors, see Figure 4.

Notably, the fit indices of these two models were not changed when predictive pathways were specified. That means the fit indices of SEM Model 2 and SEM Model 4 are the same as the original Model 2 and 4. The complete fit indices of the SEM models can be found in supplementary materials. A caveat related to the SEM results is the poor reliability scores of the direct-ideomotor and perceptual-cognitive factors (Table 3). In future studies, the same models can be tested with a bigger pool of items, which can enhance reliability scores of factors (e.g., Woody et al., 2005), in order to reconfirm the obtained results.

These results corroborate SATH’s claim that simulation-adaptation suggestions encompass the essential components of suggestibility. In other words, SATH claims that simulation-adaptation suggestions need a balanced interaction between vital top-down cognitive processes employed for responding to different forms of suggestions. Based on SATH, cognitive simulation and top-down downregulation of sensory information (i.e., sensory adaption) are both required for responding to simulation-adaptation suggestions, distinguishing them from other forms of suggestions. This proposition has a notable implication for using standardized scales of hypnotic susceptibility in clinical and experimental usage, as will be discussed below.

Regarding the SEM results, one question remains; can our results be explained by the fact that the challenge-ideomotor factor is measured by more items than the other factors? One should notice that in Model 2, both the direct-ideomotor and challenge-ideomotor factors are measured with a large number of items (4 and 5, respectively), and still, only the challenge-ideomotor factor can significantly predict the outcome of the direct-ideomotor factor, and not the other way around. This, in our opinion, is pointing that the SEM results are not due to the number of items measuring different factors or stability of factors. However, to answer this question more confidently, in future studies, one should use a bigger item pool for measuring each Factor, similar to Woody et al. (2005).

Figure 6 shows the distributions of factors’ scores (for details, see Table 1). Notably, obtained distributions of each factor were fitted to several prominent discrete distributions, that is, beta-binomial, Poisson, and negative binomial distributions. Based on calculated relative AIC and BIC, it was decided which distribution explains the data better. As shown in Table 5, all three distributions are beta-binomial. However, only the simulation-adaptation factor showed estimated α < 1 and β < 1, indicating a U-shape distribution. It should be noticed that α < 1 & β < 1, α = β = 1, and α > 1 & β > 1 represent U-shaped, uniform, and bell-shaped distributions, respectively. Therefore, the challenge-ideomotor factor can be cautiously considered distributed U-shape. By considering the structure of minor factors, the fit of the proposed models, and the distribution of factors’ scores, we can finally discuss the observed bimodal distribution of the HGSHS:A scores (see Fig. 1). Importantly, this bimodality was also observed in many other studies (for review, see Balthazard & Woody, 1989). Three possible explanations may be offered for this distribution. (1) The bimodality in HGSHS:A total scores may be the consequence of overdispersion of a beta-binomial distribution. In other words, the bimodality may be caused by smearing of the peak of the distribution and can be considered noise. (2) If we consider that the challenge-ideomotor factor has a U-shaped beta-binomial distribution and that the challenge-ideomotor factor predicts both the direct-ideomotor and cognitive-perceptual factors, we may also suggest that the bimodality in HGSHS:A is forced by the distribution of the challenge-ideomotor factor. Therefore, hypnotic suggestibility may be related to a trait with a U-shape distribution in the population, and assuming a normal distribution does not represent this facet. In other words, there is a higher chance that a person would be either fully capable or entirely incapable of responding to simulation-adaption suggestions rather than in between. Based on the discussion regarding the processes involved in simulation-adaptation suggestions, one may predict that in a given participant, the interaction between cognitive simulation and sensory adaptation either succeeds in making cognitive simulation dominant over sensory inputs, rendering this participant capable of responding to simulation-adaption (the challenge-ideomotor factor) and cognitive-simulation suggestions (direct-ideomotor), or otherwise the participant will be low-hypnotic-suggestible. Importantly, this conclusion does not mean that all hypnotic-suggestible participants are the same. That is, a given participant might have a powerful imagination (i.e., cognitive simulation) and therefore needs less sensory adaptation (i.e., top-down downregulation of sensory input), and another might be very talented in sensory adaptation, and hence, is less dependent on cognitive simulation (e.g., Terhune et al., 2011).

**Figure 6.**
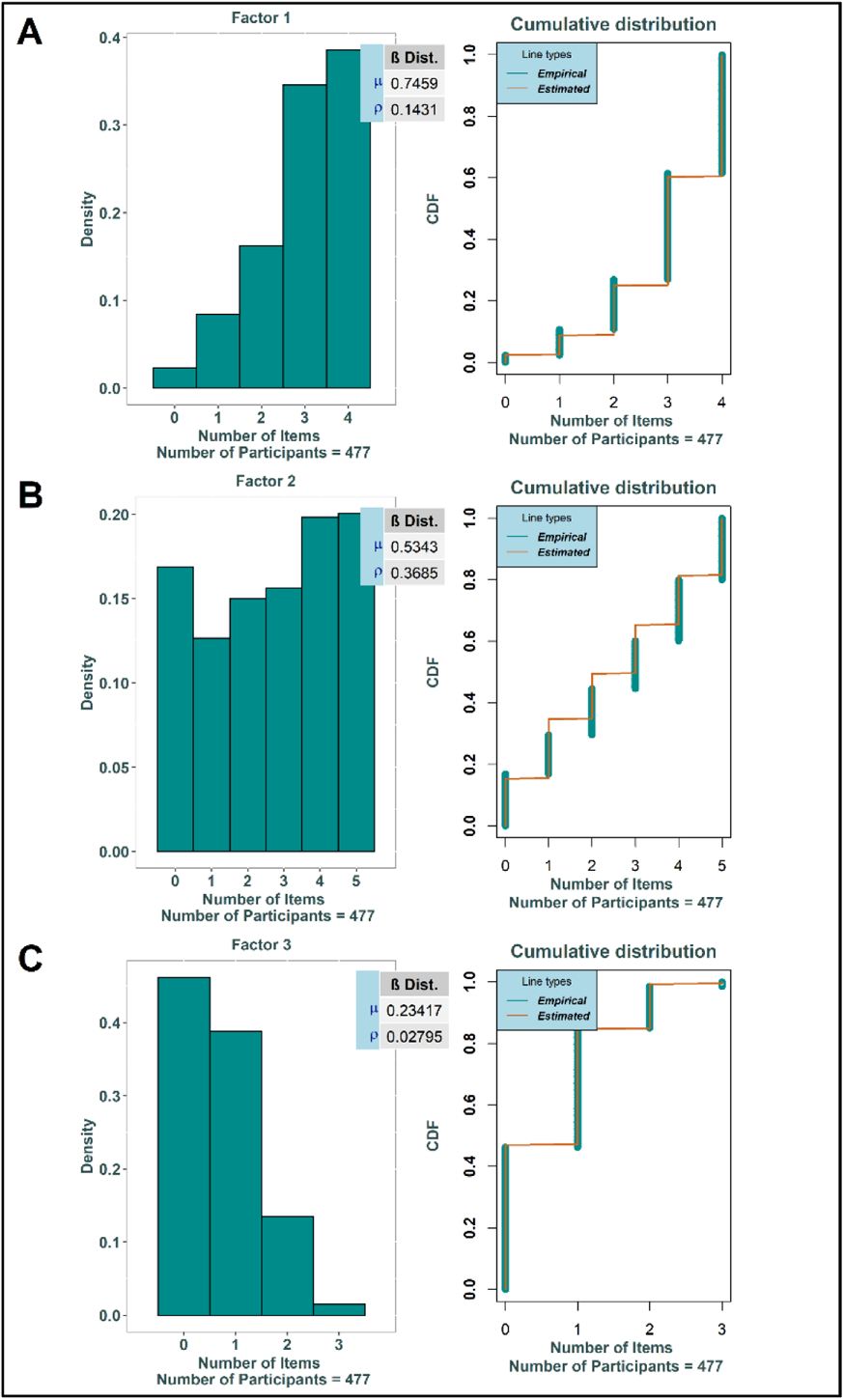
Distributions of the (A) F1: direct-ideomotor suggestions, (B) F2: challenge-ideomotor suggestions, and (C) F3: perceptual-cognitive suggestions scores. Left: Histograms of distributions. Right: Cumulative distributions of the empirical and estimated data, based on a beta-binomial distribution by the given μ and ρ.

**Table 5.** Fit indices of the distributions of the HGSHS:A scores and minor factors

| Distribution | Fitted Distribution | $\Delta AIC^a$ | $\Delta BIC^a$ | Estimated $\alpha$ and $\beta$ |
| --- | --- | --- | --- | --- |
| HGSHS:A scores | **Beta-Binomial** | **0** | **0** | **4.81 - 4.28** |
|  | Poisson | 45 | 42 | - |
|  | Negative Binomial | 47 | 48 | - |
| F2 scores | **Beta-Binomial** | **0** | **0** | **.99 - .79** |
|  | Poisson | 204 | 200 | - |
|  | Negative Binomial | 199 | 199 | - |
| F1 scores | **Beta-Binomial** | **0** | **0** | **4.46 - 1.52** |
|  | Poisson | 328 | 324 | - |
|  | Negative Binomial | 326 | 330 | - |
| F3 scores | **Beta-Binomial** | **0** | **0** | **8.14 - 26.63** |
|  | Poisson | 11 | 7 | - |
|  | Negative Binomial | 13 | 13 | - |

*Note:* The endorsed distribution is highlighted in bold.
F1: direct-ideomotor suggestions; F2: challenge-ideomotor suggestions; F3: perceptual-cognitive suggestions.
<sup>a</sup> When $\Delta AIC > 10$ , the model with minimum AIC will be endorsed (Burnham & Anderson, 2016). $\Delta AIC = AIC_i - AIC_{minimum}$ , and $\Delta BIC = BIC_i - BIC_{minimum}$ . The AIC are corrected for the finite sampling ( $AIC_c$ ).
\* $p < .05$ ; \*\* $p < .01$ ; \*\*\* $p < .001$

Finally, (3) the observed bimodality might be due to the superimposition of two unimodal distributions. Especially, given that the bifactor- (*SI*. − 1) model (Model 4) was the best fit for the data, one may assume that hypnotic suggestibility is related to two underlying mechanisms, namely, direct verbal suggestibility and hypnotizability, both of which have a normal distribution in the general population. However, as they might have positive versus negative skewness, when added together under the general concept of hypnotic susceptibility, they may cause the HGSHS:A scores to have a bimodal distribution. It must be noted that these interpretations need to be experimentally investigated before one can draw any firm conclusion. The bimodality of general hypnotic susceptibility scores is also investigated using taxometric analysis (Oakman & Woody, 1996; Reshetnikov & Terhune, 2020); however, the results are mixed regarding the existence of a high-hypnotic-suggestible subgroup (i.e., taxon). It should be noted that even if the general hypnotic susceptibility scores would not favor the existence of distinct subgroups (such as the high-hypnotic-suggestibles), one might still suggest that some of the HGSHS:A dimensions, rather than general scores, are bimodal.

## 4. General Discussion

Our results attest to the advantages of CFA and SEM over EFA. First, since CFA is theory driven and less subjected to post hoc interpretations, the results can be interpreted with greater confidence. For instance, the bifactor- (*SI*. − 1) model consists of two parts, a G-factor and the basic three-factor model. Regarding the basic three-factor part, there is an exact correspondence between the theory-driven basic three-factor model and the dominant solution of previous EFAs (e.g., McConkey et al., 1980) (see Tab. 1). However, in previous EFA (e.g., McConkey et al., 1980; Oakman & Woody, 1996; Piesbergen & Peter, 2006), minor factors and their indicators were data-driven and explained post hoc, distinguishing them from theory-driven (a priori) hypotheses needed for CFA. This issue hindered previous EFAs in agreeing on the most relevant model since several statistically-indistinguishable models fitted the data similarly well. For instance, even though some studies favor a two-factor model (e.g., Sadler & Woody, 2004), some others prefer a three- (McConkey et al., 1980) or four-factor models (e.g., Woody et al., 2005). In the current study, on the other hand, the latent factors and their indicators were derived from SATH (Zahedi & Sommer, 2021), and therefore, the uncertainty about the number and nature of latent factors were minimized. Notably, Woody et al. (2005) used CFA in order to show that a hierarchical model with a second-level G-factor and four first-level latent factors is superior to a simple multifactorial model with only four first-level latent factors; however, their CFA was focused on the possibility of adding a second-level G-factor to their EFA results and did not try to compare different models.

Second, even though multiple studies showed that the HGSHS:A scores are best explained by more than one latent factor (McConkey et al., 1980; Woody et al., 2005), to the best of our knowledge, no previous study has applied bifactor modeling to the HGSHS:A scores or any other standardized hypnotic-susceptibility scale. However, bifactor modeling is of great relevance for analyzing hypnotic suggestibility according to the proposition of Kirsch (1997) and SATH. In these accounts, standardized scales such as the SSHS:C or HGSHS:A measure hypnotic suggestibility, consisting of two components, namely, suggestibility and hypnotizability. Our results show that adding a G-factor, measuring the common variance of all items of HGSHS:A, to the basic multifactor model significantly enhanced model fit. This G-factor hypothetically measures global hypnotizability. Hence, the current results corroborate the idea that in addition to the specific minor factors, hypothetically tapping into different suggestibilities, a G-factor, arguably measuring hypnotizability, is necessary to explain the variance of the HGSHS:A scores fully.

Third, previous studies (McConkey et al., 1980; Woody et al., 2005) have demonstrated multiple hypnotic suggestibilities but did not investigate the predictive pathways between these latent factors by using SEM as their analysis was focused on factorability of hypnotic-susceptibility scores. One interpretation of our SEM results is that the outcome of the challenge-ideomotor suggestions, which hypothetically requires a combination of the two critical top-down processes involved in suggestibility (i.e., cognitive simulation and sensory adaptation), can significantly predict the outcomes of both the direct-ideomotor and cognitive-perceptual but not vice versa.

In his seminal work, Hilgard (1965b) conducted a series of EFA, investigating the structure of hypnotic susceptibility. Based on the obtained results, he proposed that responding to suggestions might depend on two “necessary but not sufficient” variables: general hypnotizability and specific cognitive processes required for responding to different groups of suggestions. Conspicuously, there is considerable overlap between the hypotheses of Hilgard (1965b) and SATH. There are two noticeable differences, however, that are worth mentioning. First, Hilgard (1965b) categorized suggestions based on their contents into six groups: agnosia and cognitive distortion, positive hallucinations, negative hallucinations, dreams and regressions, amnesia and posthypnotic suggestion, and finally, loss of motor control (Hilgard, 1965a, 1965b). Secondly, these categories were not associated with specific underlying cognitive processes, and therefore, it is not apparent why they should be distinguished. Notably, in his later works (e.g., Hilgard, 1973), Hilgard offered the neo-dissociation theory, where he postulated the existence of a hidden observer (i.e., the amnestic barrier that separates consciousness into two divisions) and connected hypnotizability to the capability of experiencing amnesia, which is different from the approach taken by SATH. However, in his earlier works, his conceptualization and distinction of general hypnotizability and specific processes are perspicuously similar to postulations of SATH. Our results have notable implications for the future applications of standardized tests in clinical and experimental settings. In line with other studies (Sadler & Woody, 2004, 2021; Woody et al., 2005), results indicate HGSHS:A is not a homogenous instrument as a single G-factor model cannot reasonably fit the data. This underscores that the HGSHS:A total scores might not be appropriate for predicting participants’ responses to experimental suggestions. This conclusion is in line with the findings of Woody et al. (2005); after conducting EFA, they used extracted factor scores and general scores for predicting participants’ responsiveness to specific suggestions in several previous studies. Their results showed that different hypnotic-susceptibility subtypes and general hypnotic-susceptibility scores differentially predict responsiveness to targeted suggestions in those experiments. However, the bifactorial structure of hypnotic-susceptibility scores highlights the necessity of employing not only simple subscale scores but also hypnotic-susceptibility profiles, which contain estimates of both G-factor scores. Additionally, the Rasch analysis results, in line with other studies (e.g., Acunzo & Terhune, 2021; McConkey et al., 1980; Sadler & Woody, 2004, 2021; Woody & Barnier, 2008), confirmed that suggestions related to different minor factors are not homogenous in terms of difficulty, which can bias subscale scores that are derived from these instruments. Therefore, it is advisable to adjust or develop standardized tests that have more balanced items measuring each minor factor.

Regarding the problem-solving suggestions, we mentioned that SATH assumes any suggestion can be a problem-solving, depending on several factors, such as the context in which the suggestion is being presented and its wording. This assumption comes from two sets of observations; first, Woody et al. (2005) observed that the arm rigidity and arm immobilization suggestions in SHSS:C load on the “perceptual-cognitive” factor which is not the case in HGSHS:A. They discussed that this difference between SHSS:C and HGSHS:A loadings is due to different contexts in which these suggestions are presented (i.e., previous suggestions). Further, it is observed that similar suggestions with different wordings can elicit different experiences in participants. For example, goal-directed imaginative compared to non-imaginative suggestions (Spanos, 1971), direct compared to indirect suggestions (Lynn et al., 1987), and elaborated-indirect compared to short-direct suggestions (Matthews et al., 1985) cause a higher sense of involuntariness or passivity. Therefore, in line with these findings, SATH hypothesizes that wording and context of suggestions can affect how participants try to approach them.

Further, SATH makes another assumption that needs to be assessed by future studies. That is, whether the cognitive processes that are used for responding to suggestions inside and outside of hypnosis are similar. There are several studies that show there is a high correlation between responding to suggestions inside and outside of hypnosis (e.g., Braffman & Kirsch, 1999; Ludwig et al., 2014; Mazzoni et al., 2009; McGeown et al., 2012; Parris & Dienes, 2013); however, based on these observations one can only tentatively assume that these correlations are due the similar cognitive processes being used inside and outside of hypnosis for responding to suggestions. Before one can firmly confirm this assumption, it needs to be tested rigorously.

It is conspicuous that there are many similarities between models proposed by SATH and previous models derived from EFA. For instance, if one disregards the hypothesis of SATH regarding the existence of two sources of variability and only focuses on the cognitive processes suggested by SATH, it is possible to propose a simple multifactorial model based on SATH; this model would be very similar or even identical to some EFA solutions (e.g., McConkey et al., 1980; Woody et al., 2005). In our opinion, this is a positive point for SATH, as it can propose a correct multifactorial model. However, the bifactor model proposed by SATH is not a hierarchical-multifactorial model, and the G-factor in this model has a remarkably different interpretation, which is already discussed in length in the results and discussion section.

We intentionally used conventional words for referring to latent factors to prevent confusion; that is, “direct-ideomotor”, “challenge-ideomotor”, and “perceptual-cognitive” (e.g., McConkey et al., 1980; Woody et al., 2005). It is appreciable that these names originate from studies conducted by Eysenck (1943) and Eysenck and Furneaux (1945), which investigated general suggestibility. However, one should notice that these names categorically dissociate movement and non-movement suggestions, which is not in complete accordance with previous results. For instance, in EFA conducted by Woody et al. (2005), it is found that the fly suggestion in HGSHS:A, and to a lesser extent, the mosquito hallucination in the SHSS:C, also loads on the “direct-ideomotor” factor, and simultaneously, the arm-rigidity suggestion in SHSS:C loads mainly on the “perceptual-cognitive” factor. These findings highlight the issue of dissociating suggestions based on being motor-related instead of using underlying cognitive processes for categorizing them.

Several limitations of our study must be discussed. First, non-significant loadings on some factors are a common anomaly in bifactor modeling (Eid et al., 2017). We also observed this issue in a conventional bifactor model of the HGSHS:A scores (see supplementary materials). Therefore, based on the suggestions of Eid et al. (2018) and Eid et al. (2017), we used the bifactor- (*S* − 1) and (*SI*. − 1) models instead of the conventional bifactor model. Even though the anomalies were better controlled in these models, some indicators still had nonsignificant loadings (see Fig. 4). Even though these anomalies were interpreted post hoc (see section 3.2. Confirmatory Factor Analyses), they were not expected at the time of postulating these models.

Further, in our study, we only employed HGSHS:A for modeling hypnotic-suggestibility scores. In future studies, however, it will be beneficial to (1) use a bigger pool of items (for instance, by using multiple standardized hypnotic-susceptibility scales, as done by Woody et al. (2005)), (2) and further measure the hypothesized top-down cognitive functions along with hypnotic-susceptibility scores, such as top-down sensory adaptation and cognitive simulation (that are postulated by SATH to be important in responding to suggestions). However, one should notice that for measuring some of these top-down cognitive functions, there is no standardized test available, and therefore, novel instruments have to be developed first. For instance, sensory adaptation is commonly investigated by using a one-item test (e.g., Lopresti-Goodman et al., 2013). Despite its relevance in basic studies, using a one-item test to gauge individual differences might result in misleading findings, as it does not account for measurement errors or task-specific variances.

Noticeably, in the current study, we tested a hypothesis derived from SATH and not SATH itself. That means, the current study cannot directly show that the cognitive processes hypothesized by SATH are being used by participants; however, if one, based on SATH, assumes that there are specific cognitive and psychosocial processes required for responding to suggestions, then it can be expected that our hypothesized bifactor model will fit the data better than hierarchical models. The current study investigated this specific hypothesis, and the obtained results are corroborating it.

In conclusion, in the current study, we conducted a CFA of HGSHS:A. Going beyond the results of previous EFAs (e.g., McConkey et al., 1980; Woody et al., 2005), our results show that (1) HGSHS:A is best explained by a bifactor model, in which a G-factor is assumed beside three correlated minor factors. These results highlight that two sources of variability affect hypnotic-susceptibility scores independently, which is of utmost importance for understanding hypnotizability as a general trait. (2) SEM of latent factors revealed that the outcome of challenge-ideomotor suggestions could predict the outcome of other suggestions. Therefore, challenge-ideomotor suggestions might require more fundamental mechanisms than the other two groups of suggestions. (3) Finally, our results have substantial implications for future applications of the standardized scales of hypnotic susceptibility. First, in line with the conclusions of Acunzo and Terhune (2021), the current study shows the need for developing new scales of hypnotic suggestibility, which can provide unbiased subscale scores. Second, in line with the proposition of Woody et al. (2005) and Jensen et al. (2017), the current results show the importance of using suggestibility profiles rather than general scores as inclusion criteria for experimental studies.

## Acknowledgment

We thank David B. Terhune, John F. Kihlstrom, and two other unknown reviewers for their detailed assessment and enlightening comments, which enhanced the quality of the manuscript. This work was supported by a scholarship to A.Z. by the Deutscher Akademischer Austauschdienst (DAAD).

## Data and Supplementary Materials

For accessing raw data, model descriptions, and R-codes, please see: https://osf.io/dus9e (DOI 10.17605/OSF.IO/DUS9E)

## Footnotes

1 Even though, it is possible that different CFA with distinguishable sets of hypotheses propose a similar model (i.e., differently interpret the model), but as CFA require a priori hypotheses regarding loadings of indicators on factors and relationships between factors, their interpretations still would be, even though different, a priori.

2 More familiar indices of beta binomial distribution can be calculated by □=□ (1−□)/□ and □=(1−□)(1−□)/□.

3 When χ^2^ test is not significant the model fits the data.

## References

1. Acunzo, D. J., & Terhune, D. B. (2021). A Critical Review of Standardized Measures of Hypnotic Suggestibility. International Journal of Clinical and Experimental Hypnosis, 69(1), 50–71. https://doi.org/10.1080/00207144.2021.1833209

2. Adams, R. A., Shipp, S., & Friston, K. J. (2013). Predictions not commands: active inference in the motor system. Brain Structure & Function, 218(3), 611–643. https://doi.org/10.1007/s00429-012-0475-5

3. Andersen, E. B. (1973). A goodness of fit test for the rasch model. Psychometrika, 38(1), 123–140. https://doi.org/10.1007/bf02291180

4. Balthazard, C. G., & Woody, E. Z. (1989). Bimodality, dimensionality, and the notion of hypnotic types. International Journal of Clinical and Experimental Hypnosis, 37(1), 70–89. https://doi.org/10.1080/00207148908410534

5. Barber, T. X. (1999). A comprehensive three-dimensional theory of hypnosis.

6. Bongartz, W. (1985). German norms for the Harvard Group Scale of Hypnotic Susceptibility, Form A. International Journal of Clinical and Experimental Hypnosis, 33(2), 131–139. https://doi.org/10.1080/00207148508406643

7. Braffman, W., & Kirsch, I. (1999). Imaginative suggestibility and hypnotizability: an empirical analysis. Journal of Personality and Social Psychology, 77(3), 578–587. https://doi.org/10.1037//0022-3514.77.3.578

8. Brown, H., Adams, R. A., Parees, I., Edwards, M., & Friston, K. (2013). Active inference, sensory attenuation and illusions. Cognitive Processing, 14(4), 411–427. https://doi.org/10.1007/s10339-013-0571-3

9. Brown, R. J., & Oakley, D. A. (2004). An integrative cognitive theory of hypnosis and high hypnotizability. The highly hypnotizable person: Theoretical, experimental and clinical issues, 152–186.

10. Burnham, K. P., & Anderson, D. R. (2016). Multimodel Inference. Sociological Methods & Research, 33(2), 261–304. https://doi.org/10.1177/0049124104268644

11. Clark, A. (2013). Whatever next? Predictive brains, situated agents, and the future of cognitive science. Behavioral and Brain Sciences, 36(3), 181–204. https://doi.org/10.1017/S0140525X12000477

12. Cohen, J. (2016). Statistical Power Analysis. Current Directions in Psychological Science, 1(3), 98–101. https://doi.org/10.1111/1467-8721.ep10768783

13. Cooper, S. R., Jackson, J. J., Barch, D. M., & Braver, T. S. (2019). Neuroimaging of individual differences: A latent variable modeling perspective. Neurosci Biobehav Rev, 98, 29–46. https://doi.org/10.1016/j.neubiorev.2018.12.022

14. Coulacoglou, C., & Saklofske, D. H. (2017). Advances in Latent Variable Measurement Modeling. In Psychometrics and Psychological Assessment (pp. 67–88). https://doi.org/10.1016/b978-0-12-802219-1.00004-3

15. Delignette-Muller, M. L., & Dutang, C. (2015). fitdistrplus: AnRPackage for Fitting Distributions. Journal of Statistical Software, 64(4). https://doi.org/10.18637/jss.v064.i04

16. Eid, M., Geiser, C., Koch, T., & Heene, M. (2017). Anomalous results in G-factor models: Explanations and alternatives. Psychological Methods, 22(3), 541–562. https://doi.org/10.1037/met0000083

17. Eid, M., Krumm, S., Koch, T., & Schulze, J. (2018). Bifactor Models for Predicting Criteria by General and Specific Factors: Problems of Nonidentifiability and Alternative Solutions. J Intell, 6(3). https://doi.org/10.3390/jintelligence6030042

18. Elkins, G. R., Barabasz, A. F., Council, J. R., & Spiegel, D. (2015). Advancing research and practice: The revised APA Division 30 definition of hypnosis. American Journal of Clinical Hypnosis, 57(4), 378–385. https://doi.org/10.1080/00029157.2015.1011465

19. Eysenck, H. J. (1943). Suggestibility and Hysteria. Journal of Neurology and Psychiatry, 6(1-2), 22–31. https://doi.org/10.1136/jnnp.6.1-2.22

20. Eysenck, H. J., & Furneaux, W. D. (1945). Primary and secondary suggestibility; an experimental and statistical study. Journal of Experimental Psychology, 35, 485–503. https://doi.org/10.1037/h0054976

21. Farah, M. J. (1988). Is visual imagery really visual? Overlooked evidence from neuropsychology. Psychological Review, 95(3), 307–317. https://doi.org/10.1037/0033-295x.95.3.307

22. Frank, T. D. (2016). Perception adapts via top-down regulation to task repetition: A Lotka-Volterra-Haken modeling analysis of experimental data. Journal of Integrative Neuroscience, 15(1), 67–79. https://doi.org/10.1142/S0219635216500059

23. Friston, K. (2010). The free-energy principle: a unified brain theory? Nature Reviews: Neuroscience, 11(2), 127–138. https://doi.org/10.1038/nrn2787

24. Galea, V., Woody, E. Z., Szechtman, H., & Pierrynowski, M. R. (2010). Motion in response to the hypnotic suggestion of arm rigidity: a window on underlying mechanisms. International Journal of Clinical and Experimental Hypnosis, 58(3), 251–268. https://doi.org/10.1080/00207141003760561

25. Green, J. P., & Lynn, S. J. (2011). Hypnotic responsiveness: expectancy, attitudes, fantasy proneness, absorption, and gender. International Journal of Clinical and Experimental Hypnosis, 59(1), 103–121. https://doi.org/10.1080/00207144.2011.522914

26. Hammond, D. C. (1998). Hypnotic induction and suggestion. American Society of Clinical Hypnosis.

27. Harrington, D. (2009). Confirmatory factor analysis. Oxford university press.

28. Hesslow, G. (2002). Conscious thought as simulation of behaviour and perception. Trends in Cognitive Sciences, 6(6), 242–247. https://doi.org/10.1016/s1364-6613(02)01913-7

29. Hilgard, E. R. (1965a). Hypnosis. Annual Review of Psychology, 16, 157–180. https://doi.org/10.1146/annurev.ps.16.020165.001105

30. Hilgard, E. R. (1965b). Hypnotic susceptibility.

31. Hilgard, E. R. (1973). A neodissociation interpretation of pain reduction in hypnosis. Psychological Review, 80(5), 396–411. https://doi.org/10.1037/h0020073

32. Jamieson, G. A. (2016). A unified theory of hypnosis and meditation states: The interoceptive predictive coding approach.

33. Jensen, M. P., Adachi, T., Tome-Pires, C., Lee, J., Osman, Z. J., & Miro, J. (2015). Mechanisms of hypnosis: toward the development of a biopsychosocial model. International Journal of Clinical and Experimental Hypnosis, 63(1), 34–75. https://doi.org/10.1080/00207144.2014.961875

34. Jensen, M. P., Jamieson, G. A., Lutz, A., Mazzoni, G., McGeown, W. J., Santarcangelo, E. L., … Terhune, D. B. (2017). New directions in hypnosis research: strategies for advancing the cognitive and clinical neuroscience of hypnosis. Neurosci Conscious, 3(1). https://doi.org/10.1093/nc/nix004

35. Jones, B., & Spanos, N. P. (1982). Suggestions for altered auditory sensitivity, the negative subject effect and hypnotic susceptibility: A signal detection analysis. Journal of Personality and Social Psychology, 43(3), 637–647. https://doi.org/10.1037/0022-3514.43.3.637

36. Jung, R. E., Flores, R. A., & Hunter, D. (2016). A New Measure of Imagination Ability: Anatomical Brain Imaging Correlates. Frontiers in Psychology, 7, 496. https://doi.org/10.3389/fpsyg.2016.00496

37. Kihlstrom, J. F. (2008). The domain of hypnosis, revisited. The Oxford handbook of hypnosis: Theory, research and practice, 21–52.

38. Kihlstrom, J. F., & Register, P. A. (1984). Optimal scoring of amnesia on the Harvard Group Scale of Hypnotic Susceptibility, Form A. International Journal of Clinical and Experimental Hypnosis, 32(1), 51–57. https://doi.org/10.1080/00207148408416000

39. Kirsch, I. (1997). Suggestibility or hypnosis: what do our scales really measure? International Journal of Clinical and Experimental Hypnosis, 45(3), 212–225. https://doi.org/10.1080/00207149708416124

40. Kirsch, I., & Lynn, S. J. (1997). Hypnotic involuntariness and the automaticity of everyday life. American Journal of Clinical Hypnosis, 40(1), 329–348. https://doi.org/10.1080/00029157.1997.10403402

41. Kirsch, I., & Lynn, S. J. (1998). Dissociation theories of hypnosis. Psychological Bulletin, 123(1), 100–115. https://doi.org/10.1037/0033-2909.123.1.100

42. Landry, M., Lifshitz, M., & Raz, A. (2017). Brain correlates of hypnosis: A systematic review and meta-analytic exploration. Neurosci Biobehav Rev, 81(Pt A), 75–98. https://doi.org/10.1016/j.neubiorev.2017.02.020

43. Li, C. H. (2016). Confirmatory factor analysis with ordinal data: Comparing robust maximum likelihood and diagonally weighted least squares. Behavior Research Methods, 48(3), 936–949. https://doi.org/10.3758/s13428-015-0619-7

44. Lopresti-Goodman, S. M., Turvey, M. T., & Frank, T. D. (2013). Negative hysteresis in the behavioral dynamics of the affordance “graspable”. Atten Percept Psychophys, 75(5), 1075–1091. https://doi.org/10.3758/s13414-013-0437-x

45. Ludwig, V. U., Stelzel, C., Krutiak, H., Magrabi, A., Steimke, R., Paschke, L. M., … Walter, H. (2014). The suggestible brain: posthypnotic effects on value-based decision-making. Social Cognitive and Affective Neuroscience, 9(9), 1281–1288. https://doi.org/10.1093/scan/nst110

46. Lynn, S. J., & Green, J. P. (2011). The sociocognitive and dissociation theories of hypnosis: toward a rapprochement. International Journal of Clinical and Experimental Hypnosis, 59(3), 277–293. https://doi.org/10.1080/00207144.2011.570652

47. Lynn, S. J., Green, J. P., Polizzi, C. P., Ellenberg, S., Gautam, A., & Aksen, D. (2019). Hypnosis, hypnotic phenomena, and hypnotic responsiveness: Clinical and Research Foundations-A 40-Year Perspective. International Journal of Clinical and Experimental Hypnosis, 67(4), 475–511. https://doi.org/10.1080/00207144.2019.1649541

48. Lynn, S. J., Laurence, J. R., & Kirsch, I. (2015). Hypnosis, suggestion, and suggestibility: an integrative model. American Journal of Clinical Hypnosis, 57(3), 314–329. https://doi.org/10.1080/00029157.2014.976783

49. Lynn, S. J., Neufeld, V., & Matyi, C. L. (1987). Inductions versus suggestions: Effects of direct and indirect wording on hypnotic responding and experience. Journal of Abnormal Psychology, 96(1), 76–79. https://doi.org/10.1037/0021-843x.96.1.76

50. Lynn, S. J., Rhue, J. W., & Weekes, J. R. (1990). Hypnotic involuntariness: A social cognitive analysis. Psychological Review, 97(2), 169–184. https://doi.org/10.1037/0033-295x.97.2.169

51. Mair, P., & Hatzinger, R. (2007). Extended Rasch Modeling: TheeRmPackage for the Application of IRT Models inR. Journal of Statistical Software, 20(9). https://doi.org/10.18637/jss.v020.i09

52. Martin, J. R., & Pacherie, E. (2019). Alterations of agency in hypnosis: A new predictive coding model. Psychological Review, 126(1), 133–152. https://doi.org/10.1037/rev0000134

53. Matthews, W. J., Bennett, H., Bean, W., & Gallagher, M. (1985). Indirect versus direct hypnotic suggestions--an initial investigation: a brief communication. International Journal of Clinical and Experimental Hypnosis, 33(3), 219–223. https://doi.org/10.1080/00207148508406650

54. Mazzoni, G., Rotriquenz, E., Carvalho, C., Vannucci, M., Roberts, K., & Kirsch, I. (2009). Suggested visual hallucinations in and out of hypnosis. Consciousness and Cognition, 18(2), 494–499. https://doi.org/10.1016/j.concog.2009.02.002

55. McConkey, K. M., Sheehan, P. W., & Law, H. G. (1980). Structural analysis of the Harvard Group Scale of Hypnotic Susceptibility, Form A. International Journal of Clinical and Experimental Hypnosis, 28(2), 164–175. https://doi.org/10.1080/00207148008409838

56. McGeown, W. J., Venneri, A., Kirsch, I., Nocetti, L., Roberts, K., Foan, L., & Mazzoni, G. (2012). Suggested visual hallucination without hypnosis enhances activity in visual areas of the brain. Consciousness and Cognition, 21(1), 100–116. https://doi.org/10.1016/j.concog.2011.10.015

57. Musek, J. (2017). The Overall Strength of the GFP. In The General Factor of Personality (pp. 183–202). https://doi.org/10.1016/b978-0-12-811209-0.00007-8

58. Näring, G. W. B., Hoogduin, K. A. L., & Keijser, C. M. P. (2004). A Rasch Analysis of the Stanford Hypnotic Susceptibility Scale, Form C. International Journal of Clinical and Experimental Hypnosis, 52(3), 250–259. https://doi.org/10.1080/0020714049052350

59. Norman, D. A., & Shallice, T. (1986). Attention to Action. In Consciousness and Self-Regulation (pp. 1–18). https://doi.org/10.1007/978-1-4757-0629-1_1

60. Oakley, D. A., Walsh, E., Mehta, M. A., Halligan, P. W., & Deeley, Q. (2021). Direct verbal suggestibility: Measurement and significance. Consciousness and Cognition, 89, 103036. https://doi.org/10.1016/j.concog.2020.103036

61. Oakman, J. M., & Woody, E. Z. (1996). A taxometric analysis of hypnotic susceptibility. Journal of Personality and Social Psychology, 71(5), 980–991. https://doi.org/10.1037/0022-3514.71.5.980

62. Parris, B. A., & Dienes, Z. (2013). Hypnotic suggestibility predicts the magnitude of the imaginative word blindness suggestion effect in a non-hypnotic context. Consciousness and Cognition, 22(3), 868–874. https://doi.org/10.1016/j.concog.2013.05.009

63. Peter, B., Vogel, S. E., Prade, T., Geiger, E., Mohl, J. C., & Piesbergen, C. (2014). Hypnotizability, personality style, and attachment: an exploratory study, part 1-general results. American Journal of Clinical Hypnosis, 57(1), 13–40. https://doi.org/10.1080/00029157.2014.906152

64. Piccione, C., Hilgard, E. R., & Zimbardo, P. G. (1989). On the degree of stability of measured hypnotizability over a 25-year period. Journal of Personality and Social Psychology, 56(2), 289–295. https://www.ncbi.nlm.nih.gov/pubmed/2926631

65. Piesbergen, C., & Peter, B. (2006). An investigation of the factor structure of the Harvard Group Scale of Hypnotic Susceptibility, Form A (HGSHS:A). Contemporary Hypnosis, 23(2), 59–71. https://doi.org/10.1002/ch.311

66. Polczyk, R. (2016). Factor structure of suggestibility revisited: new evidence for direct and indirect suggestibility. Current Issues in Personality Psychology, 2, 87–96. https://doi.org/10.5114/cipp.2016.60249

67. Preacher, K. J., & Coffman, D. L. (2006). Computing power and minimum sample size for RMSEA. In.

68. R Core Team. (2013). R: A language and environment for statistical computing.

69. Reise, S. P. (2012). The rediscovery of bifactor measurement models. Multivariate behavioral research, 47(5), 667–696.

70. Reshetnikov, M., & Terhune, D. (2020). Taxometric evidence for a dimensional latent structure of hypnotic suggestibility. https://doi.org/10.31234/osf.io/8j9va

71. Robin, B. R., Kumar, V. K., & Pekala, R. J. (2005). Direct and indirect scales of hypnotic susceptibility: resistance to therapy and psychometric comparability. International Journal of Clinical and Experimental Hypnosis, 53(2), 135–147. https://doi.org/10.1080/00207140590927617

72. Rosseel, Y. (2012). lavaan: AnRPackage for Structural Equation Modeling. Journal of Statistical Software, 48(2). https://doi.org/10.18637/jss.v048.i02

73. Sadler, P., & Woody, E. Z. (2004). Four decades of group hypnosis scales: what does item-response theory tell us about what we’ve been measuring? International Journal of Clinical and Experimental Hypnosis, 52(2), 132–158. https://doi.org/10.1076/iceh.52.2.132.28092

74. Sadler, P., & Woody, E. Z. (2021). Multicomponent Theories of Hypnotizability: History and Prospects. International Journal of Clinical and Experimental Hypnosis, 69(1), 27–49. https://doi.org/10.1080/00207144.2021.1833210

75. Shor, R. E. (1962). Three dimensions of hypnotic depth. International Journal of Clinical and Experimental Hypnosis, 10, 23–38. https://doi.org/10.1080/00207146208415862

76. Shor, R. E., & Orne, E. C. (1962). Harvard group scale of hypnotic susceptibility. Consulting Psychologists Press.

77. Shor, R. E., & Orne, E. C. (1963). Norms on the Harvard group scale of hypnotic susceptibility, form A. International Journal of Clinical and Experimental Hypnosis, 11, 39–47. https://doi.org/10.1080/00207146308409226

78. Spanos, N. P. (1971). Goal-directed fantasy and the performance of hypnotic test suggestions. Psychiatry, 34(1), 86–96. https://doi.org/10.1080/00332747.1971.11023658

79. Spanos, N. P., Cobb, P. C., & Gorassini, D. R. (1985). Failing to resist hypnotic test suggestions: a strategy for self-presenting as deeply hypnotized. Psychiatry, 48(3), 282–292. https://doi.org/10.1080/00332747.1985.11024288

80. Tanaka, J. S. (1987). “How Big Is Big Enough?”: Sample Size and Goodness of Fit in Structural Equation Models with Latent Variables. Child Development, 58(1). https://doi.org/10.2307/1130296

81. Terhune, D. B. (2015). Discrete response patterns in the upper range of hypnotic suggestibility: A latent profile analysis. Consciousness and Cognition, 33, 334–341. https://doi.org/10.1016/j.concog.2015.01.018

82. Terhune, D. B., Cardena, E., & Lindgren, M. (2011). Dissociative tendencies and individual differences in high hypnotic suggestibility. Cognitive Neuropsychiatry, 16(2), 113–135. https://doi.org/10.1080/13546805.2010.503048

83. Terhune, D. B., Cleeremans, A., Raz, A., & Lynn, S. J. (2017). Hypnosis and top-down regulation of consciousness. Neurosci Biobehav Rev, 81(Pt A), 59–74. https://doi.org/10.1016/j.neubiorev.2017.02.002

84. Terhune, D. B., & Hedman, L. R. A. (2017). Metacognition of agency is reduced in high hypnotic suggestibility. Cognition, 168, 176–181. https://doi.org/10.1016/j.cognition.2017.06.026

85. Terhune, D. B., & Oakley, D. A. (2020). Hypnosis and Imagination. In. Cambridge University Press. https://doi.org/10.1017/9781108580298.043

86. Varga, K., Farkas, L., & Mero, L. (2012). On the objectivity of the scoring of Harvard Group Scale of Hypnotic Susceptibility. International Journal of Clinical and Experimental Hypnosis, 60(4), 458–479. https://doi.org/10.1080/00207144.2012.675298

87. Weitzenhoffer, A. M., & Hilgard, E. R. (1962). Stanford hypnotic susceptibility scale, form C (Vol. 27). Consulting Psychologists Press.

88. Woody, E. Z., & Barnier, A. J. (2008). Hypnosis scales for the twenty-first century: What do we need and how should we use them. The Oxford handbook of hypnosis: Theory, research, and practice, 255–282.

89. Woody, E. Z., Barnier, A. J., & McConkey, K. M. (2005). Multiple hypnotizabilities: differentiating the building blocks of hypnotic response. Psychological Assessment, 17(2), 200–211. https://doi.org/10.1037/1040-3590.17.2.200

90. Woody, E. Z., & McConkey, K. M. (2003). What we don’t know about the brain and hypnosis, but need to: a view from the Buckhorn Inn. International Journal of Clinical and Experimental Hypnosis, 51(3), 309–338. https://doi.org/10.1076/iceh.51.3.309.15523

91. Xia, Y., & Yang, Y. (2019). RMSEA, CFI, and TLI in structural equation modeling with ordered categorical data: The story they tell depends on the estimation methods. Behavior Research Methods, 51(1), 409–428. https://doi.org/10.3758/s13428-018-1055-2

92. Yee, T. W. (2010). TheVGAMPackage for Categorical Data Analysis. Journal of Statistical Software, 32(10). https://doi.org/10.18637/jss.v032.i10

93. Yee, T. W. (2015). Vector generalized linear and additive models: with an implementation in R. Springer.

94. Zahedi, A., Abdel Rahman, R., Sturmer, B., & Sommer, W. (2019). Common and specific loci of Stroop effects in vocal and manual tasks, revealed by event-related brain potentials and posthypnotic suggestions. Journal of Experimental Psychology: General, 148(9), 1575–1594. https://doi.org/10.1037/xge0000574

95. Zahedi, A., Luczak, A., & Sommer, W. (2020). Modification of food preferences by posthypnotic suggestions: An event-related brain potential study. Appetite, 151, 104713. https://doi.org/10.1016/j.appet.2020.104713

96. Zahedi, A., & Sommer, W. (2021). How hypnotic suggestions work – critical review of prominent theories and a novel synthesis. https://doi.org/10.31234/osf.io/mp9bs

97. Zahedi, A., Stuermer, B., Hatami, J., Rostami, R., & Sommer, W. (2017). Eliminating stroop effects with post-hypnotic instructions: Brain mechanisms inferred from EEG. Neuropsychologia, 96, 70–77. https://doi.org/10.1016/j.neuropsychologia.2017.01.006

98. Zahedi, A., Sturmer, B., & Sommer, W. (2020). Can posthypnotic suggestions boost updating in working memory? Behavioral and ERP evidence. Neuropsychologia, 148, 107632. https://doi.org/10.1016/j.neuropsychologia.2020.107632

99. Zumbo, B. D., Gadermann, A. M., & Zeisser, C. (2007). Ordinal Versions of Coefficients Alpha and Theta for Likert Rating Scales. Journal of Modern Applied Statistical Methods, 6(1), 21–29. https://doi.org/10.22237/jmasm/1177992180

100. Zumbo, B. D., & Kroc, E. (2019). A Measurement Is a Choice and Stevens’ Scales of Measurement Do Not Help Make It: A Response to Chalmers. Educational and Psychological Measurement, 79(6), 1184–1197. https://doi.org/10.1177/0013164419844305

