## Supplementary Material for "Can Hypnotic Susceptibility be Explained by Bifactor Models? Structural Equation Modeling of the Harvard Group Scale of Hypnotic Susceptibility - Form A"

Author Note

* Corresponding Author; Institut für Psychologie, Humboldt-Universität zu Berlin, Rudower Chaussee 18, 12489, Berlin, Germany.

Acknowledgment

This work was supported by a scholarship to A.Z. by the Deutscher Akademischer Austauschdienst (DAAD).

Data and Supplementary Materials

For accessing raw data, model descriptions, and R-codes, please see:

DOI 10.17605/OSF.IO/DUS9E

### Supplementary Materials

#### S.1. All Models assessed by confirmatory factor analysis (CFA)

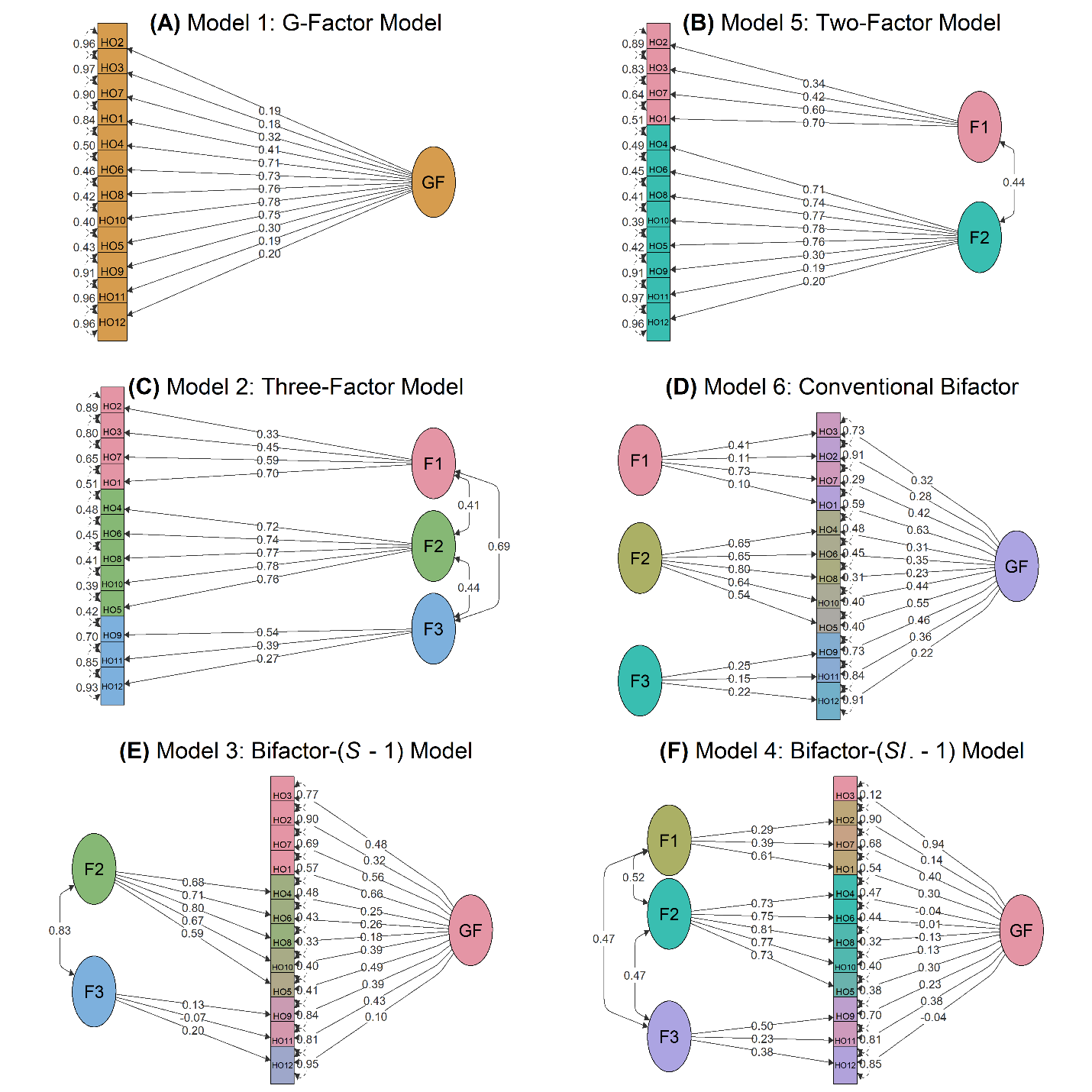

*Figure S.1.* Estimated loadings of the CFA models. (A) G-factor model (#1), (B) two-factor model (#5), (C) three-factor model (#2), (D) conventional bi-factor model (#6), (E) bifactor- ($S-1$) model (#3), and (F) bifactor- ($S.I -1$) model (#4). On the single-headed arrows, standardized factor loadings are given. All loadings in the G-factor, two-factor, and three-factor models are significant, $p < .05$. In the bifactor- $(S - 1)$ model (Model 3), the loadings of Factor 3 and HO12 on GF are not significant; however, other loadings are significant, $p < .05$. In the bifactor- $(SI. - 1)$ model (Model 4), the loadings of HO2, HO4, HO6, HO8, HO10, and HO12 on the G-factor are not significant; however, other loadings are significant, $p < .05$. In the conventional bifactor model (Model 6), the loadings of Factor 1 and 3 and HO12 on the GF are not significant; however, the other loadings are significant. Loadings are equivalent to standardized regression coefficients (beta weights), and they are estimated with diagonally weighted least squares. The self-loops show error terms. Squaring these terms gives an estimate of the variance for each item that is not accounted for by the latent construct. The curved, double-headed arrows indicating correlation coefficients between latent variables, all of which, except the correlation between F3 and F1 in bifactor- $(S- 1)$model (Model 3), are significant, $p < .05$.

*Note:* F1: cognitive-simulation suggestions; F2: simulation-adaptation suggestions; F3: problem-solving suggestions.

**Table S.1.** Fit indices of the models assessed by confirmatory factor analysis (CFA)

| Model | *df* | *χ*^2^ *^a^* | RMSEA [90% CI] *^b^* | SRMR *^c^* | CFI *^d^* | TLI *^d^* |
| --- | --- | --- | --- | --- | --- | --- |
| Model 1: G-factor model | 54 | 169.93 ^***^ | 0.067 [0.056-0.079] | 0.105 | 0.887 | 0.861 |
| Model 5: Two-factor model | 53 | 122.11 ^***^ | 0.053 [0.040-0.065] | 0.090 | 0.932 | 0.916 |
| Model 2: Three-factor model | 51 | 99.06 ^***^ | 0.045 [0.031-0.058] | 0.079 | 0.953 | 0.939 |
| Model 6: Conventional bifactor model | 42 | 77.64 ^**^ | 0.042 [0.027-0.057] | 0.068 | 0.965 | 0.945 |
| Model 3: Bifactor- ($S -1$) model | 45 | 80.95 ^**^ | 0.041 [0.026-0.055] | 0.068 | 0.965 | 0.948 |
| **Model 4: Bifactor- (**$\boldsymbol{S}\boldsymbol{.}\boldsymbol{I}\boldsymbol{-}\boldsymbol{1}$**) model** | **40** | **50.7** | **0.024 [0.000-0.042]** | **0.048** | **0.99** | **0.983** |

*Note:* The endorsed model is indicated in bold. SRMR: standardized root-mean-squared residual; RMSEA: Root Mean Square Error of Approximation; CFI: Bentler’s Comparative Fit Index; TLI: Tucker-Lewis Index.

*^a^* When *χ*^2^ test is not significant the model fits the data. However, as the $N=477$ was very large, it is expected that *H_0_* would be over-rejected.

*^b^* Lower values of RMSEA indicate better fit, with values < .05 indicating a close fit to the data (Xia & Yang, 2019). For $1-\beta>0.9$, $H0<0.05$, and $H1>0.1$ the required sample size is $N>120$.

*^c^* Lower values of SRMR indicate better fit, with SRMR < .08 indicating a close

fit to the data.

*^d^* Values > .95 for CFI and TLI indicate a good fit. TLI is not normalized and may have values > 1.

* *p* < .05; ** *p* < .01; *** *p* < .001

**Table S.2.** Results of likelihood ratio tests comparing all models

| Competing models | | *df* | *χ*^2^ *^a^* | |
| --- | --- | --- | --- | --- |
| Model 5: Two-factor model | Model 1: G-factor model | 1 | | 31.3 ^***^ |
| Model 2: Three-factor model | Model 1: G-factor model | 3 | | 47.9 ^***^ |
| Model 2: Three-factor model | Model 5: Two-factor model | 2 | | 16.8 ^***^ |
| Model 3: Bifactor- ($S -1$) | Model 1: G-factor model | 9 | | 69.6 ^***^ |
| Model 3: Bifactor- ($S -1$) | Model 5: Two-factor model | 8 | | 35.9 ^***^ |
| Model 3: Bifactor- ($S -1$) | Model 3: Three-factor model | 6 | | 18.9 ^**^ |
| Model 6: Conventional bifactor | Model 1: G-factor model | 12 | | 80.9 ^***^ |
| Model 6: Conventional bifactor | Model 5: Two-factor model | 11 | | 40.9 ^***^ |
| Model 6: Conventional bifactor | Model 2: Three-factor model | 9 | | 21.4 ^*^ |
| Model 6: Conventional bifactor | Model 3: Bifactor- ($S -1$) | 3 | | 1.9 |
| Model 4: Bifactor- ($S.I -1$) | Model 1: G-factor model | 14 | | 95.2 ^***^ |
| Model 4: Bifactor- ($S.I -1$) | Model 5: Two-factor model | 13 | | 59.2 ^***^ |
| Model 4: Bifactor- ($S.I -1$) | Model 2: Three-factor model | 11 | | 41.3 ^***^ |
| Model 4: Bifactor- ($S.I -1$) | Model 3: Bifactor- ($S -1$) | 5 | | 22.2 ^***^ |
| Model 4: Bifactor- ($S.I -1$) | Model 6: Conventional bifactor | 2 | | 20.5 ^***^ |

*Note:* *^a^* If the *χ*^2^ test is significant the more complex model will be endorsed, if not, the simpler model is endorsed.

** *p* < .01; *** *p* < .001

**Table S.3.** Fit Indices of the models assessed by structural equation modeling (SEM)

| Model | *df* | *χ*^2^ *^a^* | RMSEA [90% CI] *^b^* | SRMR *^c^* | CFI *^d^* | TLI *^d^* |
| --- | --- | --- | --- | --- | --- | --- |
| Model 2 (SEM): Three-factor model | 51 | 99.06 ^***^ | 0.045 [0.031-0.058] | 0.079 | 0.953 | 0.939 |
| Model 4 (SEM): Bifactor- ($S.I -1$) model | 40 | 50.7 | 0.024 [0.000-0.042] | 0.048 | 0.99 | 0.983 |

*Note:* The endorsed model is indicated in bold. SRMR: standardized root-mean-squared residual; RMSEA: Root Mean Square Error of Approximation; CFI: Bentler’s Comparative Fit Index; TLI: Tucker-Lewis Index.

*^a^* When *χ*^2^ test is not significant the model fits the data. However, as the $N=477$ was very large, it is expected that *H_0_* would be over-rejected.

*^b^* Lower values of RMSEA indicate better fit, with values < .05 indicating a close fit to the data (Xia & Yang, 2019). For $1-\beta>0.9$, $H0<0.05$, and $H1>0.1$ the required sample size is $N>120$.

*^c^* Lower values of SRMR indicate better fit, with SRMR < .08 indicating a close

fit to the data.

*^d^* Values > .95 for CFI and TLI indicate a good fit. TLI is not normalized and may have values > 1.

* *p* < .05; ** *p* < .01; *** *p* < .001
